# How does the response of egocentric boundary cells depend upon the coordinate system of environmental features?

**DOI:** 10.1101/2025.02.03.636060

**Authors:** Jennifer C. Robinson, Patrick A. LaChance, Samantha L. Malmberg, Mahir Patel, Elaina Gross, Danielle E. Everett, Shruthi Sankaranarayanan, Jiarui Fang, Michael E. Hasselmo

**Author notes:** Corresponding authors: Prof. Michael Hasselmo, twitter: @HasselmoMichael, Dr. Jennifer Robinson, Dr. Patrick A. LaChance.

## Abstract

Neurons in the retrosplenial (RSC) (Alexander et al 2020a, LaChance & Hasselmo 2024) and postrhinal cortex (POR) respond to environmental boundaries and configurations using egocentric coordinates relative to an animal’s current position. Neurons in these structures and adjacent structures also respond to spatial dimensions of self-motion such as running velocity (Carstensen et al 2021, Robinson et al 2023). Experimental and modeling data suggest that these responses could be essential for guiding behaviors such as obstacle avoidance and goal-directed navigation (Erdem & Hasselmo 2012, Erdem & Hasselmo 2014). However, these findings still leave the unanswered question: What specific features, and in what coordinate frames, drive these egocentric neural responses? Here we present models of the potential circuit mechanisms generating egocentric responses in RSC. One model posits that neurons encode internal representations of barriers in head-centered coordinates, defined by distance and angle, which are modulated by running velocity to enable trajectory planning and obstacle avoidance. We contrast this with a complementary hypothesis in which neurons respond to retinotopic features—such as the top, bottom, or edges of walls, which may serve as precursors to head-centered representations. Additional hypotheses include trajectory-based forward scanning (e.g., ray tracing) for barrier detection or comparing optic flow across the visual field. These hypotheses generate complementary modeling predictions about how changes in environmental parameters could alter the neural responses of egocentric boundary cells that are presented here.

## INTRODUCTION

The generation of behavior in spatial environments requires the formation of internal representations of relevant spatial features to guide behavior, to avoid obstacles and barriers, while planning and following a pathway to a desired goal location. We commonly map the position of barriers, objects, and animals relative to the world, using allocentric world-centered coordinates such as north-south, east-west (Figure 1A). However, sensory receptors on an animal initially detect environmental features in egocentric head-centered coordinates, relative to the position of the animal’s sensory receptors, such as those of the visual system (Figure 1B). In addition, actions necessary to avoid barriers need to be generated in egocentric coordinates. Thus, it is logical that there should be an internal representation of the world in egocentric coordinates, which codes the position of objects and barriers, or the center of the environment, relative to an animal (see Figure 1B).

**Figure 1.**
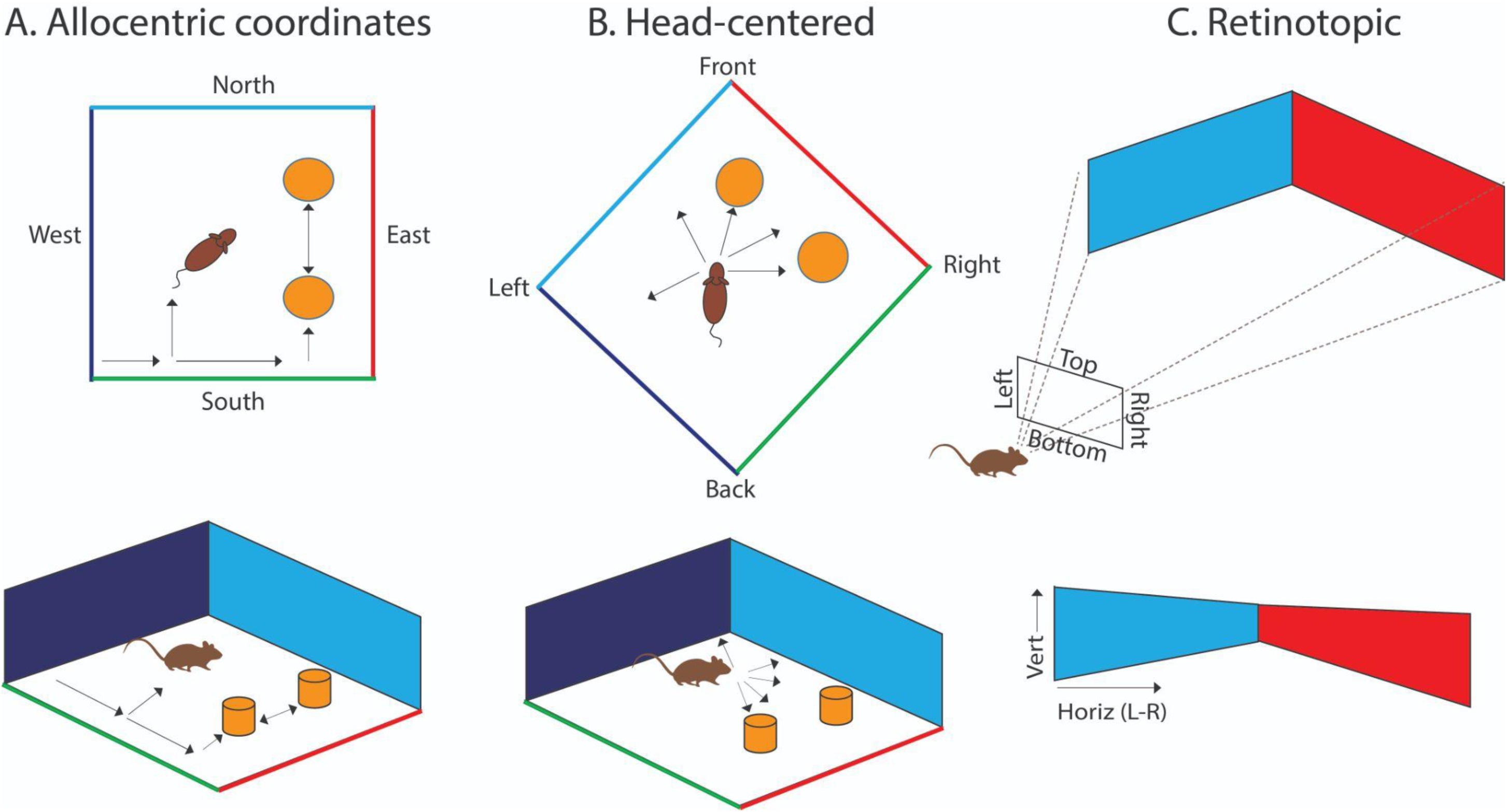
Overview of allocentric, head-centered and retinotopic coordinates. A. Allocentric coordinates. Most computer animations create a world in allocentric coordinates, defining the position of barriers, objects, and animals relative to the world coordinates of north-south, east-west. Arrows indicate position of animal and objects relative to the world. B. Head-centered coordinates. For the specific position and pose of an animal, the allocentric coordinates of barriers and objects can be rotated and translated to egocentric head-centered coordinates (front, back, left, right), based on the angle and distance relative to the animal (arrows). These head-centered coordinates determine the viewpoint of the world. C. Animals receive visual input in retinotopic coordinates that can be simulated by a projection of head-centered features onto an image plane (left-right, top-bottom). Visual features are coded in retinotopic coordinates of horizontal (left-right) and vertical (top-bottom). The simulations presented here focus on taking neural representations in retinotopic or head-centered coordinates and simulating the responses of egocentric boundary cells.

Recent neurophysiological data has demonstrated neurons that respond to environmental boundaries in egocentric coordinates. Neurons known as Egocentric Boundary Cells (EBCs) in the retrosplenial cortex (RSC) respond when environmental boundaries are at a specific angle and distance in egocentric coordinates (Alexander et al 2020a, van Wijngaarden et al 2020) (Figure 2). The models presented in this paper will focus on understanding the responses of these neurons. Similar neurons coding egocentric responses to boundaries have been shown in the dorsomedial striatum (Hinman et al 2019) and postrhinal cortex (Gofman et al 2019). Studies also show egocentric coding of the egocentric angle (egocentric bearing) and egocentric distance to the center of the environment in the postrhinal cortex (POR) and the medial entorhinal cortex (MEC) (LaChance & Taube 2023, LaChance et al 2019, Wang et al 2020, Wang et al 2018). Examples of the neural response of egocentric boundary cells are shown in Figure 2.

**Figure 2.**
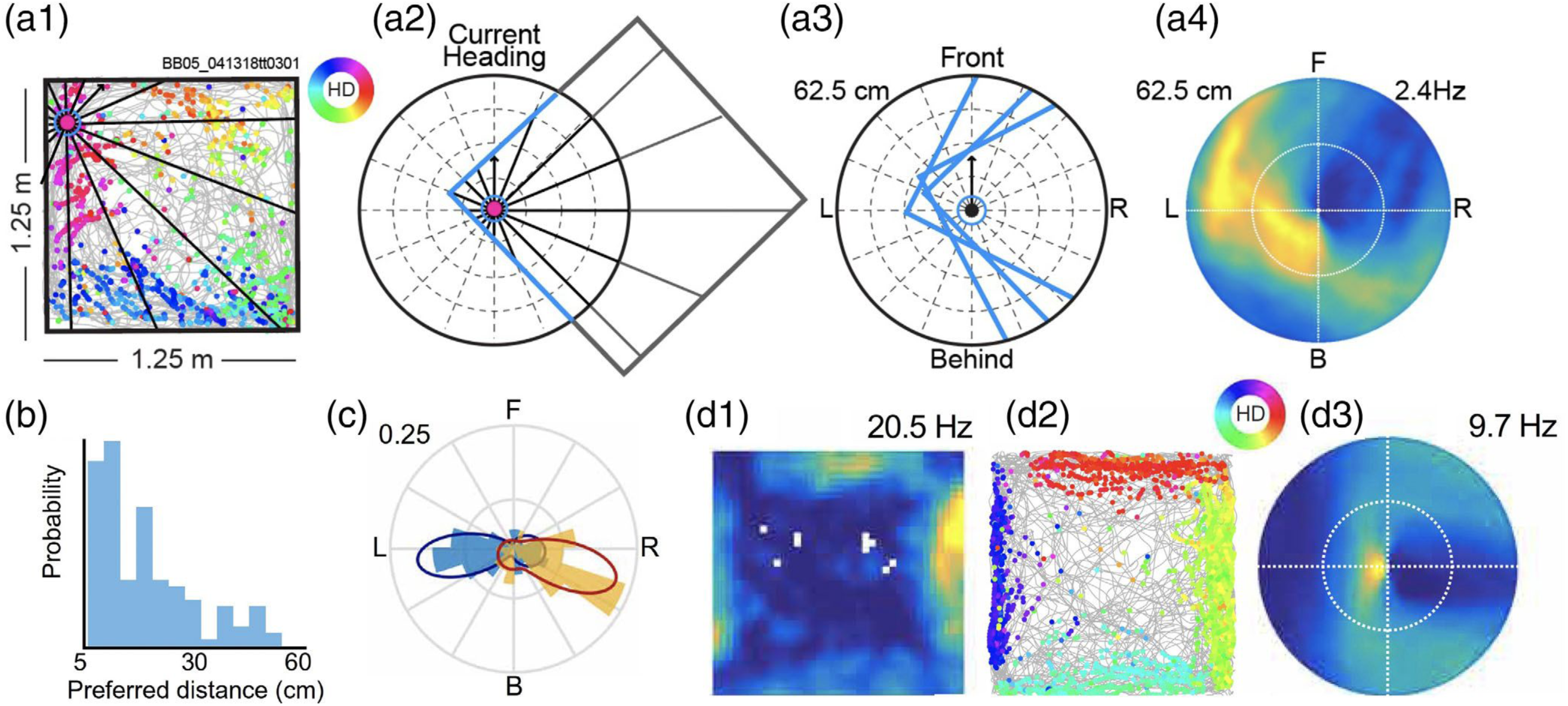
Responses of an egocentric boundary cell in retrosplenial cortex. A1. Plot in allocentric coordinates shows foraging trajectory in gray. Each time a spike occurs, animal position is plotted as a point with color indicating head direction (color wheel indicates directional coding). A2: Boundaries within 62.5 cm of animal (blue) are plotted in egocentric coordinates relative to animal head direction for a single spike. A3: Boundary positions for three spikes. A4: Example egocentric boundary ratemap constructed after repeating steps A1-A3 for all spikes and normalizing by occupation. Color axis indicates zero (blue) to peak firing (yellow) for this neuron (using MATLAB Parula colormap). B. probability distribution of preferred distance from boundary of all retrosplenial cortex egocentric boundary cells. C. polar histograms and probability density estimates of preferred boundary bearing of all retrosplenial cortex EBCs. Yellow and blue bars correspond to EBCs recorded in the left and right hemisphere, respectively. D. Another example EBC recorded in the retrosplenial cortex. D1. two-dimensional ratemap shows firing rate for animal position in 1.25m^2^ square environment (blue is no firing, yellow is peak firing rate). D2. plot of trajectory in gray with color of dot indicating head direction of animal at time of spike. D3. egocentric plot showing receptive field on the left side of the animal. *Adapted from Alexander et al., 2020*.

Avoiding collisions and planning paths to goals requires selection of movements in egocentric coordinates that depend upon egocentric coding of environmental boundaries, but also on the coding of the current running velocity. Neurons have been shown to code current running speed in structures including the entorhinal cortex (Dannenberg et al 2020, Hinman et al 2016, Kropff et al 2015) and the retrosplenial cortex (Carstensen et al 2021). These codes could be important for computing whether the specific angle and speed of running velocity will collide with a barrier at that angle within a short time or distance, in order to signal to an animal when it should turn. This process might work in a more generalizable manner in the allocentric coordinate system of entorhinal grid cells or hippocampal place cells, though these cell types are not simulated here. Previous models have proposed that forward sweeps of trajectories coded by grid cells could interact with internal representations of goals and barriers to guide pathway planning (Erdem & Hasselmo 2012, Erdem & Hasselmo 2014) and recent data from the entorhinal cortex supports this framework of forward trajectory sweeps (Vollan et al 2024). One set of models presented here will focus on the coding of the distance and angle of barriers in egocentric coordinates, and we will also discuss simulations of forward sweeps for detecting potential collisions.

Environmental features can be coded in different possible egocentric coordinate frames. One possible egocentric coordinate frame could be centered on the current position and angle of the head (head-centered coordinates). Another egocentric coordinate system could be centered on the position and angle of the body (body-centered coordinates). A further egocentric coordinate frame could be centered on the retina (retinotopic coordinates). The activation of visual receptors in the retina occurs on a two-dimensional plane in retinotopic coordinates, as shown in Figure 1C. Retinotopic coordinates can be simulated by a projection transform from the distance and angle of features in head-centered coordinates.

Thus, head-centered coordinates and retinotopic coordinates provide alternate, complementary egocentric coordinate systems for driving neurophysiological responses of egocentric boundary cells. In fact, it is probably essential that both of these coordinate frames are required at different stages in the generation of egocentric boundary responses. Retinotopic coordinates appear in early visual areas, but it is not clear how much these coordinate systems are anatomically segregated or intermixed within later-stage structures.

Initial models have shown that neural circuit responses to images in retinotopic coordinates can be effective for simulating these neural responses (Lian et al 2023). However, as shown here, when the position of features in the environment is changed, the response to features in retinotopic coordinates will generate different patterns of responses compared to the coding in head-centered coordinates. The models presented here will show how neural responses in different coordinate frames would exhibit differences in response based on manipulations of the properties of the walls and barriers of an environment, including their height, their vertical tilt, and their horizontal (azimuth) orientation. The focus of this paper is to generate predictions about how neurophysiological responses at different stages of processing would differ based on whether they are coding features in head-centered coordinates or retinotopic coordinates, and how this coding of features would respond to environmental changes.

## METHODS

The models presented here generate predictions about neurophysiological responses to environmental manipulations that have not yet been published. As an initial step in developing the models, the models start with a baseline simulation of the previous neurophysiological responses of neurons in the retrosplenial cortex that were recorded from rodents (rats and mice) as they foraged for distributed food reward (crushed up Froot Loops) in an open field environment (Alexander et al 2020a) and with the insertion of objects (Carstensen et al 2021). The simulations then move beyond the manipulations in previously published data to address predictions about the proposed response to new manipulations of parameters such as height, vertical tilt, or horizontal orientation of walls and how the response depends upon the coordinate frame of internal representations.

The simulations include several different processing components that are described in the following sections, but are first briefly summarized in this paragraph. The simulations include representations of the external environment and the animal’s viewpoint. These simulations of the environment may sound complex, but they directly correspond to the elements used in computer animation or video game engines to produce an animal’s viewpoint of an environment as it moves. These very standard parts of the simulation include: 1.) The allocentric configuration of walls, inserted barriers and objects in a spatial environment in Euclidean allocentric coordinates for wall and barrier corners relative to east-west coordinates (x), and north-south coordinates (y), 2.) In some cases, the position of the walls and barriers of the spatial environment relative to the animal in head-centered cylindrical coordinates (coding distance and angle from each feature to the animal in the horizontal x,y plane, plus height z), 3.) The position of walls and barriers in head-centered Euclidean coordinates: LR=left-right, FB=front-back, UD=up-down and finally, 4.) the retinotopic visual input to the animal using a projection transform from the head-centered 3D Euclidean coordinates LR, FB, and UD into 2D retinotopic coordinates h = horizontal and v = vertical. In the model, the features of the world are coded both in retinotopic coordinates and in head-centered coordinates. (See Figures 3 and 4). Thus, these are complementary coordinate systems essential for the standard steps used in modeling visual function in computer vision as well as generating scenes in computer animation. The necessity of both of these steps in simulating the world could correspond to a similar necessity of both coordinate systems in the internal neural representation of the world.

**Figure 3.**
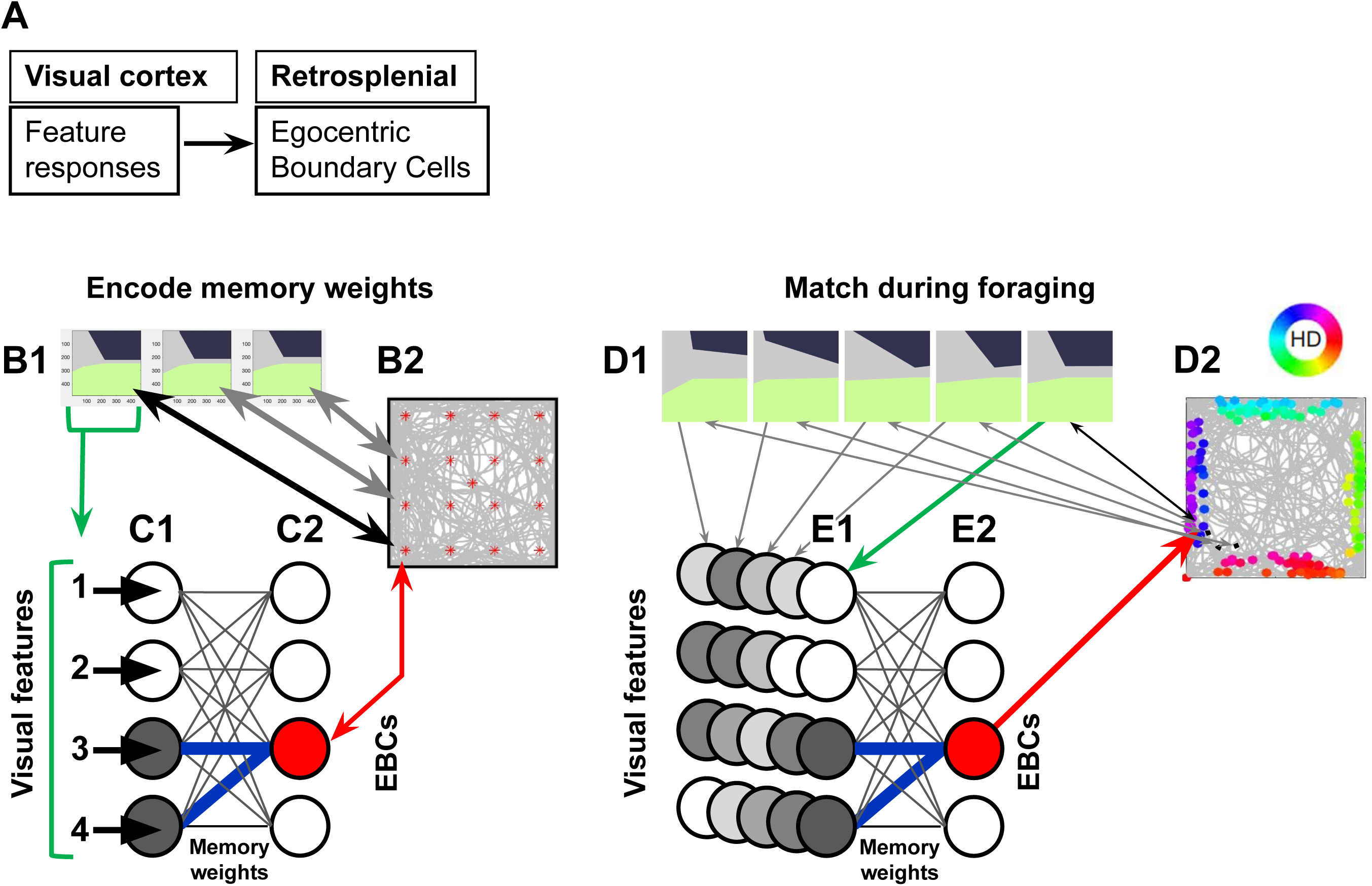
Encoding and matching in retinotopic model. A. Visual cortical areas project to retrosplenial cortex. The model simulates an array representing visual features interacting with simulations of neurons in retrosplenial cortex including egocentric boundary cells. B1. Three example retinotopic viewpoints are shown from the start of the simulation based on different agent locations show by red asterisks (see gray arrows). C. During encoding before the foraging, the 2-dimensional array of features coding individual retinotopic viewpoints (C1) are associated with activity of a single retrosplenial neuron (red) activated for that position and head direction (red arrow). Hebbian modification of synaptic weights (blue lines) creates the memory weights (template) coding the association of those features with a neuron. Lines between circles represent axons and synapses, and thickness of lines indicates strengthening of synaptic efficacy. D. At each time step during foraging, the current retinotopic visual input (D1) activates the visual feature array (E1) that then spreads across the synaptic connectivity weights determined by previous encoding. The activation of each individual neuron is computed as the match (dot product) with each row of the visual feature array with one row of the weight matrix. This activation can cross threshold and activate postsynaptic egocentric boundary cells (red circle in E2) at specific locations and head directions shown by colored dots in D2).

**Figure 4.**
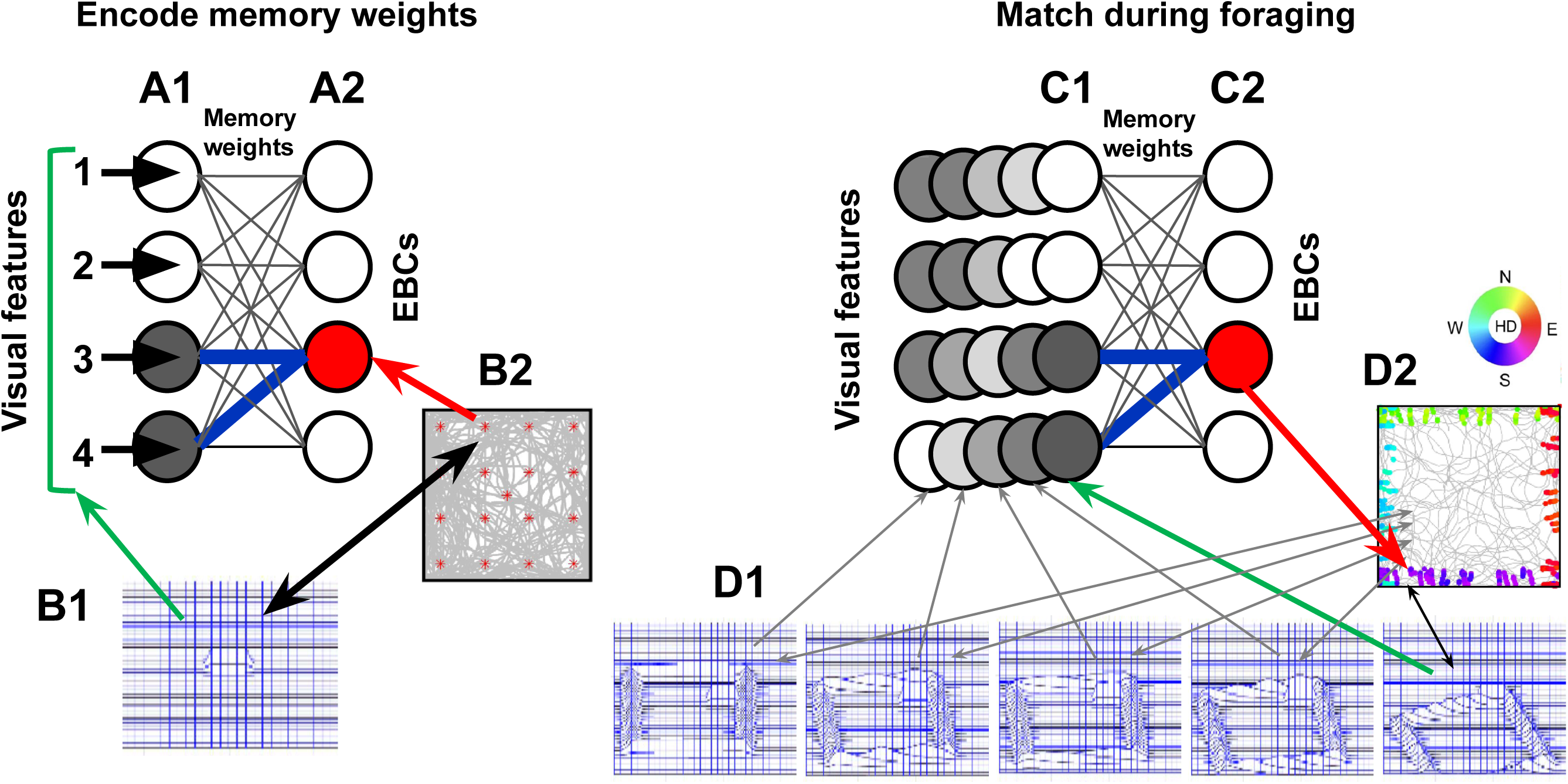
Encoding and matching in head-centered model. A1. During encoding, the head-centered model activates elements of a presynaptic vector (A1) based on the elements of an array (B1) coding sensory features of barriers based on their 3D head-centered coordinates. B1 shows array code for local barrier features in a location (B2) where barrier is right in front of the animal (black line to red asterisk). A2. For each location (red asterisks), the postsynaptic activity involves activation of one EBC (red) for the location in B2. For each location, the Hebbian rule modifies presynaptic to postsynaptic weights (blue). C1. During foraging, each time step evokes a presynaptic array corresponding to the head-centered features for all barriers viewed from the current location at that time step (D1). Note that the peak bumps in the center of each D1 plot represent the memory weights for an initially viewed wall segment, and the surrounding walls represent the current environment. The head-centered array activity spreads across synaptic weights to activate postsynaptic neurons (C2). When the features of all barriers match weights (blue) for a specific segment (from B1), this activates a specific postsynaptic cell (red) above threshold, resulting in plotting of a spike (violet) on the trajectory plot (D2). The spike is plotted with a color based on head direction. The retinotopic model uses similar input arrays and matching.

The model also includes simulation of the movements of the animal as it explores, including 4.) The animal’s forward momentum, 5.) The avoidance of obstacles, and 6.) The attraction to novel inserted barriers and objects. Similar to previous simulations (Erdem & Hasselmo 2012, Erdem & Hasselmo 2014), this uses ray-tracing algorithms from computer animation to guide possible behaviors. Ray-tracing involves computing whether and where a vector arising from the animal will intersect with the surface plane of a barrier. These properties of simulated movement might ultimately be guided by the neural model, as in previous work (Erdem & Hasselmo 2012, Erdem & Hasselmo 2014), but the current version is designed to replicate relatively realistic trajectories without using neural circuits. The use of algorithms for guiding movement is standard in programs for computer animation and video game graphics, for example in generating non-player characters that move around without external control in computer games.

The primary novel components of this paper are the simulation of the mechanism of generation of egocentric boundary cells based on retinotopic coordinates (as summarized in Figure 3) or head-centered coordinates (as summarized in Figure 4), and on the forward scanning (Erdem & Hasselmo 2012, Erdem & Hasselmo 2014) of trajectories in the environment (as shown in Figure 11). These neural responses could be generated within the same brain region, but are here simulated using separate but complementary mechanisms.

In these simulations, both of these coordinate systems are proposed to be based on modifying synaptic connections between neurons in visual cortical regions and the retrosplenial cortex (Figure 3A), to simulate how EBC responses could be driven by both of these types of coordinate systems in the retrosplenial cortex. The simulations include the possible role of internal retrosplenial representations as input for the EBC responses.

The simulations involve two main phases, an encoding phase and a matching phase. The encoding phase is shown on the left side of Figures 3 and 4 (Figures 3B and 3C; Figures 4A and 4B), and the matching phase during foraging is shown on the right side of Figures 3 and 4 (Figures 3D and 3E, Figures 4C and 4D).

7)Encoding of synaptic memory weights (Figure 3C1 and 4A1). During this phase, the agent is simulated in a discrete set of encoded head directions (facing North in the figures). For each head direction during encoding, the agent is simulated in a discrete set of pre-determined positions in the environment that might cover a range of locations, including positions near different walls. The input from the environment activates sensory representation arrays (Figure 3B1 and C1, Figure 4A1 and B1) that are activated at an array of specific positions and head directions (Figure 3B2 and Figure 4B2). These input arrays consist of an array of feature-responsive neurons in different regions of visual cortex, and possibly also neurons generating recurrent connections in retrosplenial cortex. These input arrays then provide presynaptic activity to cause Hebbian modification to strengthen synapses connecting with a postsynaptic array. For this example, strengthened synaptic weights are shown by blue lines in Figure 3C1 and Figure 4A1. The postsynaptic array contains activity of individual retrosplenial neurons activated at specific positions and head directions (the red asterisks in Figure 3B2 and Figure 4B2 indicate the array of positions for the north-facing head direction in this example).

8) Matching of sensory input with previous memory. After the initial encoding of synaptic weights based on a set of discrete positions, the responses of egocentric boundary cells are then tested during a long, foraging trajectory (Figure 3D2, Figure 4D2). In this period of the simulation, the model simulates a long, complex and continuous trajectory through the environment (gray lines in Figures 3D2 and Figure 4D2). At each time step along the trajectory, the position and head direction are used to simulate the sensory input to the retinotopic array (Figure 3D1 and Figure 4D1). The matching/comparison process involves computing the dot product of the input array with each row of the synaptic memory weights (Figure 3E1 and Figure 4C1). This simulates how presynaptic sensory cortex input of perceptual features interacts with synaptic connectivity to activate individual retrosplenial neurons that store individual templates based on the synaptic weights to their dendritic trees. For each retrosplenial neuron, a trajectory plot of spiking activity is generated. For each neuron’s trajectory plot (Figure 3D2 and Figure 4D2), if the dot product crosses a threshold, a filled circle is plotted in figures of the trajectory at that position, and the color of the filled circle in the figure plot is determined by the current head direction of the animal.

This overall neural simulation framework is used to test two potentially complementary hypotheses about the coordinate systems (head-centered or retinotopic) coded by individual egocentric boundary cells that could be intermixed in the retrosplenial cortex. These hypotheses provide insights into both the mechanism of generation of these responses, as well as their potential role in guiding behavior based on ongoing transformations of internal representations by current velocity and head direction.

The first hypothesis involves retinotopic coding and the second involves head-centered coding. To provide more detail, the first hypothesis involves: 9.) Retinotopic coordinates for visual sensory input feature arrays during both encoding (Figure 3B1) and foraging (Figure 3D1). As shown in Figures 3B1 and 3D1, these are 2-dimensional arrays that contains a set of pixels each of which represents the environmental features (green floor, gray walls) in a specific point of retinotopic coordinates (h = horizontal position and v = vertical position on the retina). Previously published simulations have been effective for generating models based on retinotopic coordinate input (Lian et al., 2023), but this did not yet address the potential intermixing of a code that is retinotopic for some egocentric boundary cells and head-centered for other egocentric boundary cells (or intermixing of the coordinates within individual neurons).

As shown in Figure 4, the second hypothesis involves: 10.) Encoding of head-centered coordinates for representing environmental features. During encoding, the head-centered coordinates of the 3D position of individual components of the environment barriers are computed as an occupancy map where the values in the coordinates indicate adjacency of a barrier segment (Figure 4B1). These head-centered coordinates are computed for an array of specific agent locations (Figure 4A2 and B2). Visual input during encoding at one location activates a head-centered presynaptic array (Figure 4A1) and allows Hebbian modification of synaptic weights (blue lines in Figure 4A1) between the presynaptic array and individual postsynaptic cells (Figure 4A2) that are activate at each location (Figure 4B2). During matching, these head-centered features are then computed based on visual input at each time step of the foraging trajectory (Figure 4D1). The head-centered features activate the presynaptic array (Figure 4C1) at each time step. The presynaptic activity spreads across the synaptic weights, and this activates the postsynaptic cells (Figure 4C2) based on the match of presynaptic activity with the pattern of weights. If an individual postsynaptic cell is activated above its firing threshold, the simulation plots a spike for this cell on the gray trajectory based on the current simulated location and with a color based on the current simulated head direction (Figure 4D2). Note that the head-centered input in Figure 4D1 is further extended by 11.) Simulation of neural transformations (rotations and translations) of the head-centered coordinates of the memory of wall and barrier positions, based on the movement velocity of the animal, to allow tracking of walls and barriers that are behind the animal or in darkness.

The following sections provide more detailed description of the components of the model.

### Simulation of the environment

The next three sections describe the visual components of the model that are simply the standard components of computer animation or video game engine graphics, addressing the steps shown in Figure 1. The MATLAB simulations used here define an environment with barriers (usually walls) defined by arrays of points in 3D allocentric coordinates. Each segment of the wall is defined by a set of four points in 3D allocentric coordinates (x = west-east, y = south-north) that later allow computation of the view of that wall segment by the animal. The position and angle of an animal are defined in the environment and allow computation of the retinotopic visual images seen when the animal has a specific allocentric location (in x = east-west, y = north-south coordinates of the environment) and a specific allocentric head direction (compass direction defined in azimuth – elevation angle is assumed to be zero). In the simulation, these points are combined in an array that includes an additional row of ones for homogeneous coordinates. The use of homogeneous coordinates allows transformation of the allocentric wall coordinates into egocentric coordinates relative to the current location and head direction using affine transformation matrices for rotation and translation.

### Rodent movement

The thin gray lines in each simulation represent the movement trajectories of the animal. This includes the location of the rodent in x, y, z coordinates (the variable called place, with agent height held constant) and a separate representation of the head direction of the rodent (the variable called *angle*) at each time point. The simulation starts with an initial position and direction and then updates the position at each time point. The movement of the rodent is simulated to replicate the distributed trajectories followed when foraging for food (Figure 3B2 and 3D2, Figures 4B2 and 4D2). This movement trajectory is created using velocity and acceleration vectors that are modified randomly at each time step, but are also guided by additional algorithms to turn away from barriers. As in previous work (Erdem & Hasselmo 2012), the barrier collision avoidance algorithm uses forward extrapolation of the current velocity vector to determine potential collisions with each of the wall segments based on standard ray tracing algorithms. These algorithms reject vectors that collide with barriers, and allow selection of trajectories that do not collide with barriers within a certain distance. Most simulations use an abstract version of barrier collision avoidance, but future simulations will attempt to relate this collision avoidance to forward trajectory scanning by neuronal activity (Erdem & Hasselmo 2012, Erdem & Hasselmo 2014).

### Simulation of current egocentric input based on viewpoint

Once the foraging trajectory has been generated that avoids the environmental barriers, this trajectory is used to generate the egocentric visual input to the animal based on its viewpoint of the coordinates defining each wall segment from the position of the animal. At each time point, the current animal position is subtracted from wall points to generate the allocentric coordinate difference dx,dy,dz. Then, for each feature (often a corner connecting wall segments or a position along a wall), the egocentric head-centered distance is computed as sqrt(dx² + dy²). Similarly, the angle of each feature (wall corner) is computed as arctan(dy/dx). This generates egocentric head-centered coordinates in cylindrical coordinates (distance and angle in the horizontal plane, plus height z). Subsequently, the egocentric head-centered cylindrical coordinates are transformed into egocentric head-centered Euclidean coordinates (left-right, front-back, up-down).

### Simulation of retinotopic coordinates

Once these egocentric input coordinates have been generated, they can be used to generate the retinotopic coordinates of wall corners using the perspective transform, as shown schematically in Figure 1C and in specific examples in Figure 3B1 and 3D1. Standard computer animation or game engine graphics would take the homogeneous egocentric Euclidean coordinates of the corners of walls and apply the perspective transform from their 3D egocentric position toward the animal’s eye position, projecting this onto a two-dimensional plane of a certain size in front of the animal (called a camera projection). Graphics will simply plot patches of color within the projected region on the two-dimensional plane. The patches match the color of the environment used in experimental work. This appears as a green ground plane, gray walls, and black background in the retinotopic images computed during encoding in Figure 3B1 and during the matching process during foraging in Figure 4D1. All of the above sections of the simulations simply replicate what is standard in computer animation and video game engine graphics. The novel simulations of neural responses are in the next sections.

### Simulation of internal representations for memory matching

Everything presented above is standard in computer animation or game engines. However, the novel feature of the current simulations concerns how the input array neurons represent features in either or both retinotopic and/or head-centered coordinates for matching with a synaptic memory template. The need for matching with a synaptic memory template is suggested by the fact that the EBC neuronal response appears to be consistent for similar features (walls) across different environments and different parts of an environment. Computer animation and game engines do not commonly require matching of a stimulus with a stored template, and if they do perform this matching, they usually compute Euclidean distances based on Euclidean coordinates of component features. It would be interesting to form a neural model that codes coordinates in a continuous manner (for example, by phase of spiking), but it is not clear how to do this in single neurons for multiple dimensions. So, we will use a more standard neural code in which firing rate represents the presence of a specific feature at a specific coordinate point in either retinotopic or head-centered coordinates.

Thus, the input arrays code environmental features using two different types of coordinate systems: 1.) retinotopic arrays, or 2.) head-centered arrays. The retinotopic array is essentially the pixel version of an image (Figure 3B1 and 3D1), in which ray tracing from the 3D position of features in the environment to an image plane is used to define the retinotopic image based on where the vector for a specific pixel will collide with the image plane (or visual receptor). Using this approach, retinotopic images are created in which the environment colors match colors used in mouse experiments, with wall barriers in light gray, and the ground plane in light green (Figure 3B1 and 3D1). The open space above the environment walls (which would involve viewing of distant shadowy walls in the environment) is coded as black on the image plane.

The head-centered array maintains a value for whether a barrier or object overlaps with the 3-dimensional location coded by an individual element of the head-centered array (Figure 4B1 and 4D1). Thus, its dimensions can be mapped to head-centered polar coordinates of distance and angle, or to the head–centered Euclidean dimensions of front–back, left–right.

### Mixture of head-centered and retinotopic coding

Note that we are specifically not stating that these head-centered or retinotopic representations are the representation across the entire brain or across even one region. Instead, we are investigating whether individual neurons are using a retinotopic or head-centered representation even within a single region, which would suggest that these stages of processing may be intermixed for generating the transformation between retinotopic and head-centered coordinates. In fact, it is possible that even individual neurons have mixed selectivity for head-centered or retinotopic coordinate systems.

### Encoding of memory templates based on prior viewpoint

As noted above, the repeated firing properties of neurons to similar egocentric cues in different locations suggests that current sensory input is being matched to an internal memory template. As shown in Figure 3A, we propose that this involves an interaction of the feature arrays from visual cortical areas with neuronal activity in the retrosplenial cortex. During an encoding phase, the model receives sensory input (Figure 3B1) that generates presynaptic activity patterns (Figure 3C1) to form memory templates based on the sensory input in different coordinate systems at a set of specific locations (and head directions) visited by the agent before the start of the foraging trajectory (red asterisks in Figure 3B2, Figure 4B2, and Figure 5Av). The model creates memories by Hebbian synaptic modification. The sensory feature arrays from the environment (Figure 3B1 and Figure 4B1) sequentially activate arrays of presynaptic neuronal activity (Figures 3C1 and 4A1) when the animal is at this series of individual locations and head directions (red asterisks in Figure 3B2 and 4B2). The presynaptic activity is associated with the sequential activation of different single retrosplenial neurons (Figure 3C2 and 4A2) at these locations. For each position and head direction, Hebbian modification of synaptic weights associates the feature array with the individual neuron (blue lines in Figure 3C and Figure 4A represent synapses strengthened by Hebbian modification).

**Figure 5.**
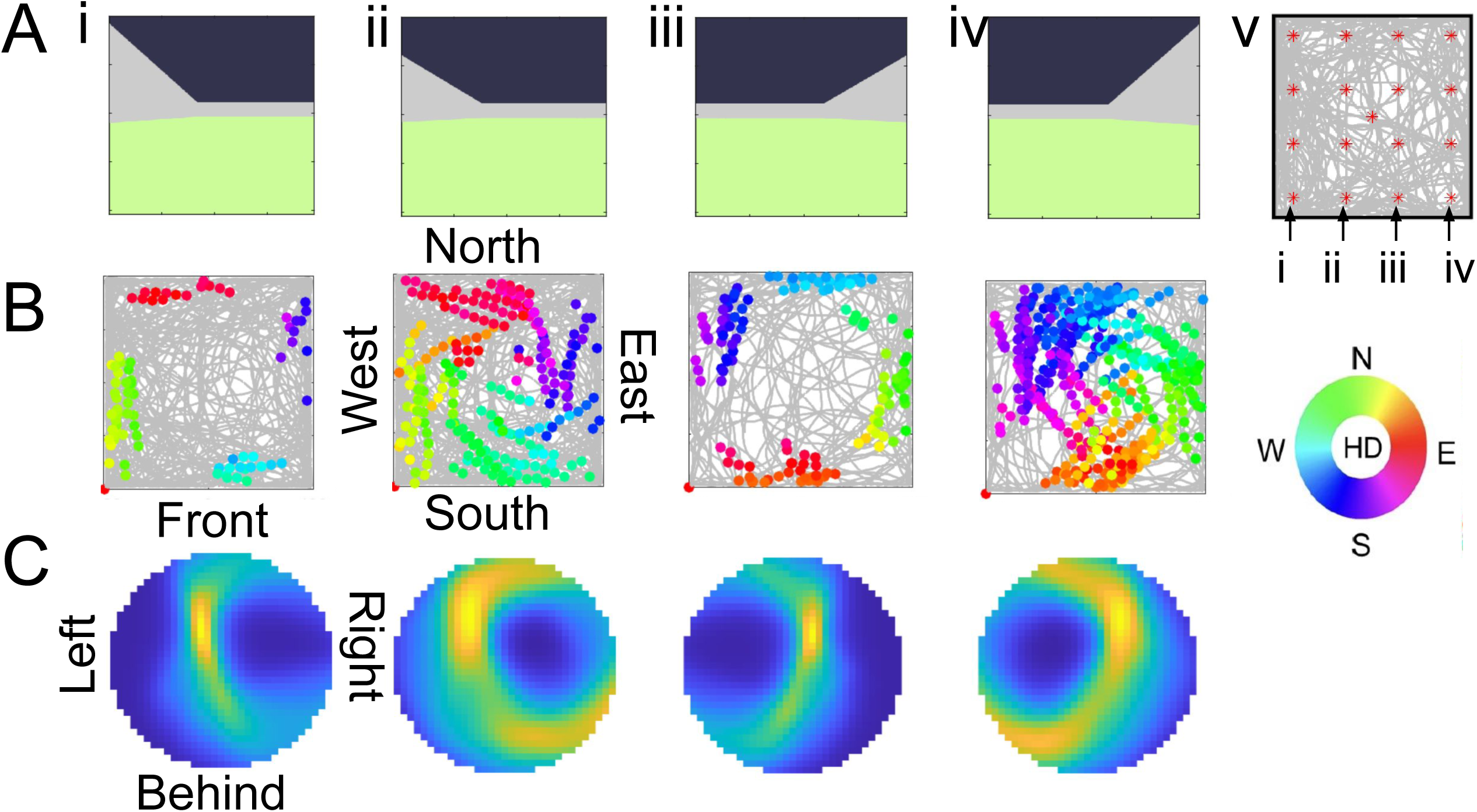
Properties of EBCs with retinotopic memory weights. A. Retinotopic images generated for four different positions at the start of the simulation to generate synaptic memory templates corresponding to the wall nearby on the left (i), further away on the left (ii), nearby on the right (iii), and further away on the right (iv). The synaptic memory weight templates then code visual features in retinotopic coordinates, and a dot product then compares these memory weight templates with the incoming sensory input array created from retinotopic coordinates at each time step along the trajectory. B. Trajectory plots showing the overall trajectory of the simulated animal in light gray, with colored dots indicating the location of the simulated animal when the dot product value for an array representing sensory responses to the synaptic memory template crosses a threshold. C. Egocentric plots averaging the egocentric polar positions containing a boundary over every example of a spike being generated. These plots show the average position of the boundary that generates a neural response of the neuron in row B of the same column, indicating responses to a wall nearby on the left (i), a wall further on the left (ii), a wall nearby on the right (iii) or a wall at a further distance on the right (iv).

For the retinotopic hypothesis, the model generates the projection transform on the image plane (Figure 3B1) at each of the grid of starting points (red asterisks in 3B2) before computing the foraging trajectory (gray line). The model then stores the retinotopic memory weights associating the initial retinotopic array of pixels (Figure 3B1) computed at the individual locations and head directions with an individual retrosplenial neuron.

The head-centered hypothesis involves a different simulation that forms the associations with the same number of retrosplenial neurons. During encoding before the start of the foraging trajectory, the simulation will form memories of initial head-centered arrays (Figure 4B1) of points at each of the grid of locations and head directions (red asterisks in Figure 4B2). For each location and head direction, the sensory input array computes a 2-dimensional array of points along the environmental barriers (in head-centered coordinates front-back, and left-right) and generates a 2-dimensional Gaussian for each of these points, so that the wall is represented by a smoothed array of points. Figure 4B1 shows the array generated during encoding when the animal is facing North near the North wall of the environment (Figure 4B2). Additional examples of encoded head-centered arrays are shown in Figure 6Ai-iv. Note that these represent walls at different angles relative to the animal, corresponding to facing north but being close to the wall in the south (6Ai, behind animal), west (6Aii, left side of animal), east (6Aiii, right side of animal), or north wall (6Aiv, in front of animal). For each of these locations, the computation process is repeated for representation of 3-dimensional representations of features at different elevations in the world (see Figure 8A), with Gaussians generated based on interpolation of that elevation relative to the top and bottom of the wall. During encoding, this head-centered input array shown in Figures 4B1 and 5Ai-iv is associated with the active postsynaptic retrosplenial neurons (Figure 4B2) by Hebbian modification of the memory weight matrix (blue lines in Figure 4B).

**Figure 6.**
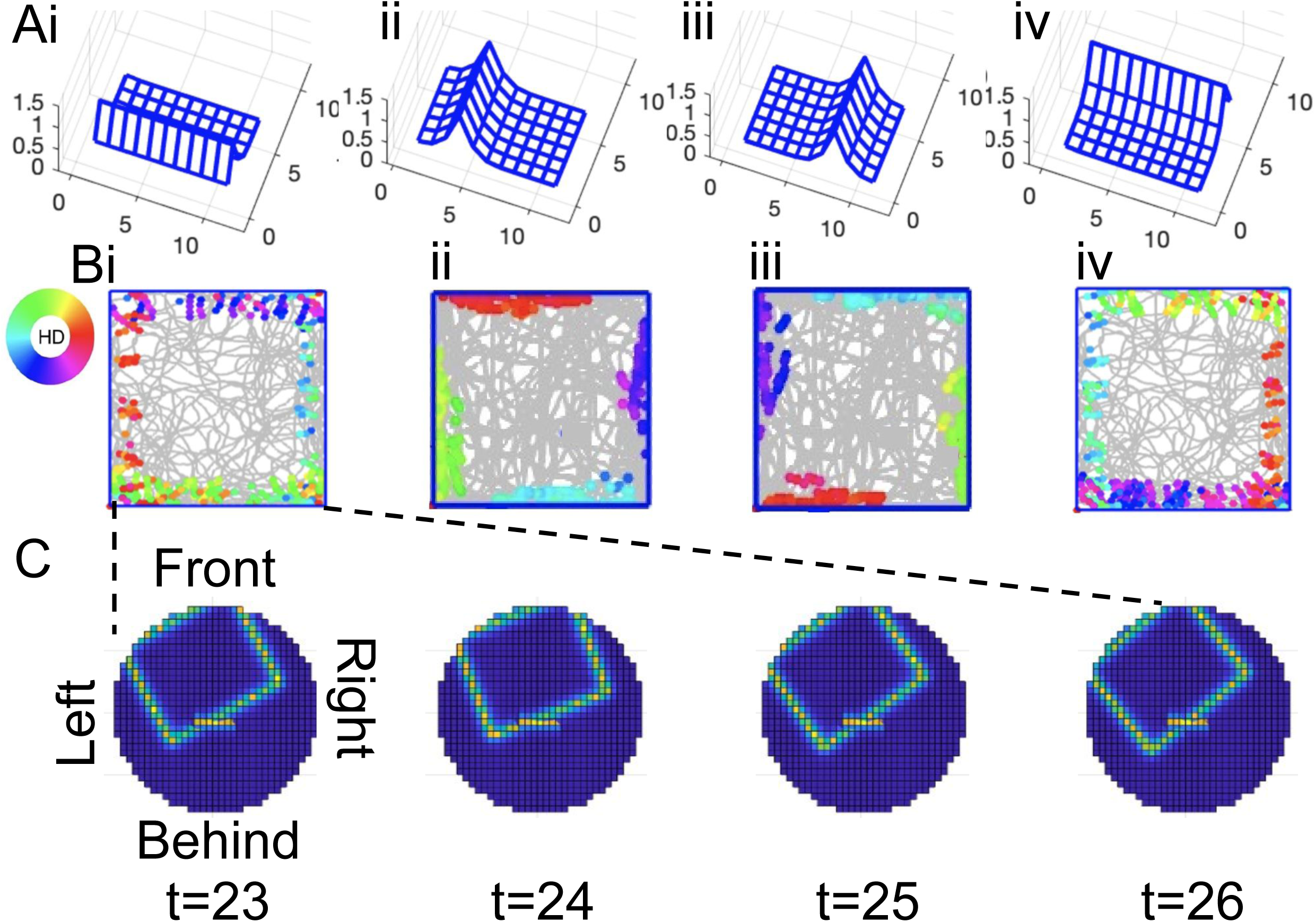
Properties of neurons with head-centered memory weights. A. Local head-centered arrays of nearby features (within 3 units of the front-back coordinate and 5 units of the left-right coordinate). These local arrays are generated for four different positions at the start of the simulation to generate synaptic memory weights corresponding to the wall behind the animal (i), on the left side (ii), on the right side (iii) and in front of the animal (iv). The memory weights code features in head-centered coordinates of distance and angle that are then coded in front-back, left-right positions. The memory weights from these arrays are compared with sensory input in head-centered coordinates at each time step along the trajectory. B. Trajectory plots showing the overall trajectory of the simulated animal in light gray, with colored dots indicating the location of the simulated animal when the dot product value for an array representing sensory responses to the synaptic memory weights crosses a threshold. C. For Figure 6Bi, an example shows four time steps of the matching/comparison process in which the local head-centered array behind the animal appears as a short horizontal line in the center of the head-centered array. The current egocentric sensory input of the barrier at each step is shown as the larger array of light colored pixels at different distances and angles. The sensory input best matches the local array at time t = 24, so would generate the largest dot product at that time point (and therefore the most spiking).

Note that we are testing the retinotopic and head-centered arrays separately, but these properties could be intermixed within the real network. Individual neurons might show signs of mixed selectivity with responses that suggest coding in both coordinate systems. Or alternately, different neurons within a single region might show coding in one or the other coordinate system.

### Matching of current sensory input with memory weights

After the encoding process described above and in Figures 3B, C and 4A, B and 5A, there is a retrieval process during the foraging trajectory. For the retinotopic model shown in Figure 3, once the foraging trajectory starts, the retinotopic sensory inputs are computed at each time point on the simulated trajectory (multiple steps of sensory input are shown in Figure 3D1). Based on the location and head direction at that time point, the current sensory input is used to generate sensory input arrays (Figure 3D1). These sensory input arrays activate a presynaptic neural array with a specific pattern of activity depending on the sensory input. Multiple time steps of presynaptic activity are illustrated in Figure 3E1. This presynaptic activity then spreads across the synaptic weight matrix. As shown in Figure 3E, this is simulated as the matrix multiplication that involves computing the dot product of the feature array with the synaptic weights in each row providing input to individual postsynaptic neurons (Figure 3E2). The computation of the dot product simulates the pattern of synaptic input matching to the pattern of synaptic strengths into the dendrites of a neuron, in which case a larger number indicates a better match to that neuron’s memory of features. If the dot product is large enough to cross a threshold the postsynaptic retrosplenial neuron will generate a spike at that specific time point (Figure 3E2). This is then plotted as a filled circle at that location along the simulated trajectory (Figure 3D2), with the color of the filled circle indicating the animal’s current head direction at that time point of the trajectory (Figure 3D2 color wheel). For the head-centered model during the matching process, the head-centered sensory input array is computed (Figure 4D1) for individual locations along a trajectory (locations indicated by gray arrows in Figure 4D2). These sensory input arrays at different positions along the trajectory sequentially activate the presynaptic array with different patterns of neural activity (Figure 4C1). These presynaptic activity patterns spread across the previously modified synaptic weight matrix to activate individual postsynaptic retrosplenial neurons (Figure 4C2). This spread of activity across the weight matrix is computed by matrix multiplication, which involves computation of a dot product between the current head-centered presynaptic array (Figure 4C1) and each row of the weight matrix to see which postsynaptic neuron is activated (Figure 4C2). When the dot product for an individual postsynaptic neuron crosses its firing threshold, then a filled circle is plotted at that time point along the foraging trajectory (gray lines in Figure 4D2) with a color corresponding to the current head direction at that point on the trajectory (color wheel indicates coding of head direction with green for North and purple for South).

## RESULTS

The model can generate simulations of egocentric boundary cell responses using the simple one step neural network. These neural networks effectively implement a matching/comparison process of input activity with the synaptic weights in each row of the matrix representing visual cortex and other cortical input to individual retrosplenial neurons. The simulations can generate egocentric boundary cells using either the retinotopic code of the horizontal and vertical position of features on the image plane, or the simulations can use the head-centered code of distance and angle to boundary features (or front-back, left-right, up-down coordinates). These hypotheses will be presented separately, but they could be intermixed in the real biological data. In our results below, the initial simulations of previous neural recordings in open field environments will be presented first. Then, predictions that have not yet been tested will be described about the effect of variations in height, vertical tilt, and horizontal orientation of barriers that will be analyzed.

### Simulations of EBC firing in an open field

The initial publications about egocentric boundary cells and egocentric bearing cells cited above primarily presented data on recordings in rectangular open field environments without any inserted objects or barriers and only a few manipulations of surrounding boundaries. The initial simulations presented here show that both the head-centered features and retinotopic feature responses can be used to generate realistic egocentric boundary cell responses in the open field environment. Thus, these previous recordings do not indicate whether the responses of the neurons are driven by retinotopic or head-centered input.

### Simulation using retinotopic features in open field

Figure 5 shows examples of EBC responses generated with memory features based on synaptic weights modified in response to features in retinotopic coordinates from the starting images shown at the top of the figure. The images reflect the fact that the current experimental environment has a green floor, gray walls, and is in a dark room surrounded by black curtains. Figure 5Ai shows an example of a retinotopic image of the wall being close to the animal on the left. Figure 5Bi shows the firing pattern of the model based on the dot product of current input features with the memory array of synaptic weights. This shows the location of the simulated agent each time the dot product value crosses a threshold to cause a spike. The different colors indicate the head direction of the simulated agent, showing that it fires near the west wall when facing north (green), fires near the north wall when facing east (red), fires near the east wall when facing south (purple) and fires near the south wall when facing West (cerulean). Figure 5Ci shows the egocentric plot relative to the simulated animal at the center of the circle, for the average boundary position over all the spikes generated by the simulated neuron. This shows that the neuron responds when a wall is close on the left side of the animal at the center. Figure 5Aii shows an image of a more distant wall on the left that was stored in the retinotopic memory template of synaptic weights. Figure 5Bii shows that the computation of a dot product generates values that cross a threshold and generate spiking at a larger distance from the wall. Figure 5Cii shows that the average wall position computed over all the spikes is at a greater distance on the left side of the animal. Figures 5A,B,Ciii show the same examples for a stored memory template using an image of a retinotopic view of a wall nearby on the right side of the animal, and figures 5A,B,Civ show this for an image of a wall at a greater distance on the right.

### Simulation using head-centered response in the open field

Figure 6 shows examples of the simulation of open field responses using feature responses based on the position of boundary cues in head-centered coordinates. The simulations shown in this section use head-centered responses to the distance and angle of each feature in the azimuth plane (i.e. analogous to the floor of the environment). That is, this is an array of sensory neurons that respond to the presence or absence of single barrier component at a specific distance and angle in head-centered coordinates (as used in previous work by Bicanski and Burgess, 2018). At each time step, the full array of neurons responding to sensory features in different head-centered positions is updated with the current locations of boundary elements.

Figure 6A shows the arrays used to create memory templates of the head-centered array at four different starting locations. For the feature response arrays shown in Figure 6A, the position of the boundary is sampled for a small set of positions around the animal, and the head-centered distance and angle of each feature is transformed into two-dimensional Euclidean coordinates, giving a small square array centered on the animal. In all examples in Figure 6A, the animal is facing to the north, and the boundary segment is in a different position relative to the animal. In Figure 6Ai, the animal is facing to the north, but located near the south wall, so the boundary segment is directly behind the animal. In Figure 6Aii, the animal is facing north in a location by the west wall, so the boundary segment is to the left of the animal. In Figure 6Aiii, the animal is facing north in a location by the east wall, so the boundary segment is to the right of the animal, and in Figure 6Aiv, the animal is facing north near the north wall boundary segment is directly in front of the animal. The encoded head-centered arrays are smaller than the actual full head-centered array of all possible positions of the boundary in the environment that is used for matching in Figure 6C. Note that the templates can be stored for a range of starting head directions, but due to the symmetry of the environment, many of the responses would be similar (i.e. near south wall facing west is similar to being near west wall facing north).

Once these head-centered memory templates are formed on the synaptic weights to the EBC neurons, then the full head-centered array is computed at each time step (Figure 6C), and the portion of this array in the smaller number of pixels around the animal spreads across the weights of the network. This results in the computation of a dot product between the current sensory input of boundary feature locations in head-centered coordinates (Figure 6C) with the memory template (Figure 6A). When this crosses a threshold, the network generates a spike shown as a colored dot on the gray trajectory in Figure 6B. The dot is colored by the head direction code shown with the color wheel in Figure 6B. Note that Figure 6Bi shows an example that would be difficult to generate with retinotopic coordinates as it uses a head-centered template behind the animal. This was encoded at a position near the south wall when the animal was facing north (Figure 6Ai). This generates neuronal responses when the animal is running away from the barrier, for example, when going south (purple) from the north wall, or going east (red) from the west wall. The other examples in Figure 6B show the spikes generated along the trajectory for a template encoding a barrier array at different positions, showing the response to a left side wall (Figure 6Bii), a right side wall (Figure 6Biii) and a wall directly in front of the animal (Figure 6Biv). Figure 6C shows the computation process for just four time steps of the trajectory in Figure 6Bi, showing a template that consists of a wall segment behind the animal, overlapped with the input of the current egocentric array of the full barrier. This shows that the head-centered barriers match best at one time point (t = 24) compared to other time points. That would yield the highest dot product and crosses the threshold to generate a spike.

### Simulations of boundary height differences during sensory input

This section describes the effect of varying the height of walls during the sensory experience of the environment, but NOT varying the height of the memory templates for each cell. The prediction of these different coordinates for the firing of EBCs will first be described for the retinotopic model and then, for the head-centered model.

### Retinotopic model response to height

Figure 7 shows examples of EBC responses generated with memory features based on synaptic weights modified in response to features in retinotopic coordinates from the starting images shown in in the left column (60 cm) of Figure 7A, B, and C. Note that the same 60 cm memory feature templates are used for all three outputs shown in the rows in Figure 7D, E, and F. Thus, Figure 7A shows that the memory template for 60 cm is used for matching with sensory input at each time step of foraging, even when the actual visual sensory input at time step has shorter walls as in the 30 cm and 15 cm examples in Figure 7A. The 30 cm and 15 cm examples in Figure 7A show how the sensory input at that same given location (12 cm from a left wall) will differ from the view used previously to create a memory at that location.

**Figure 7.**
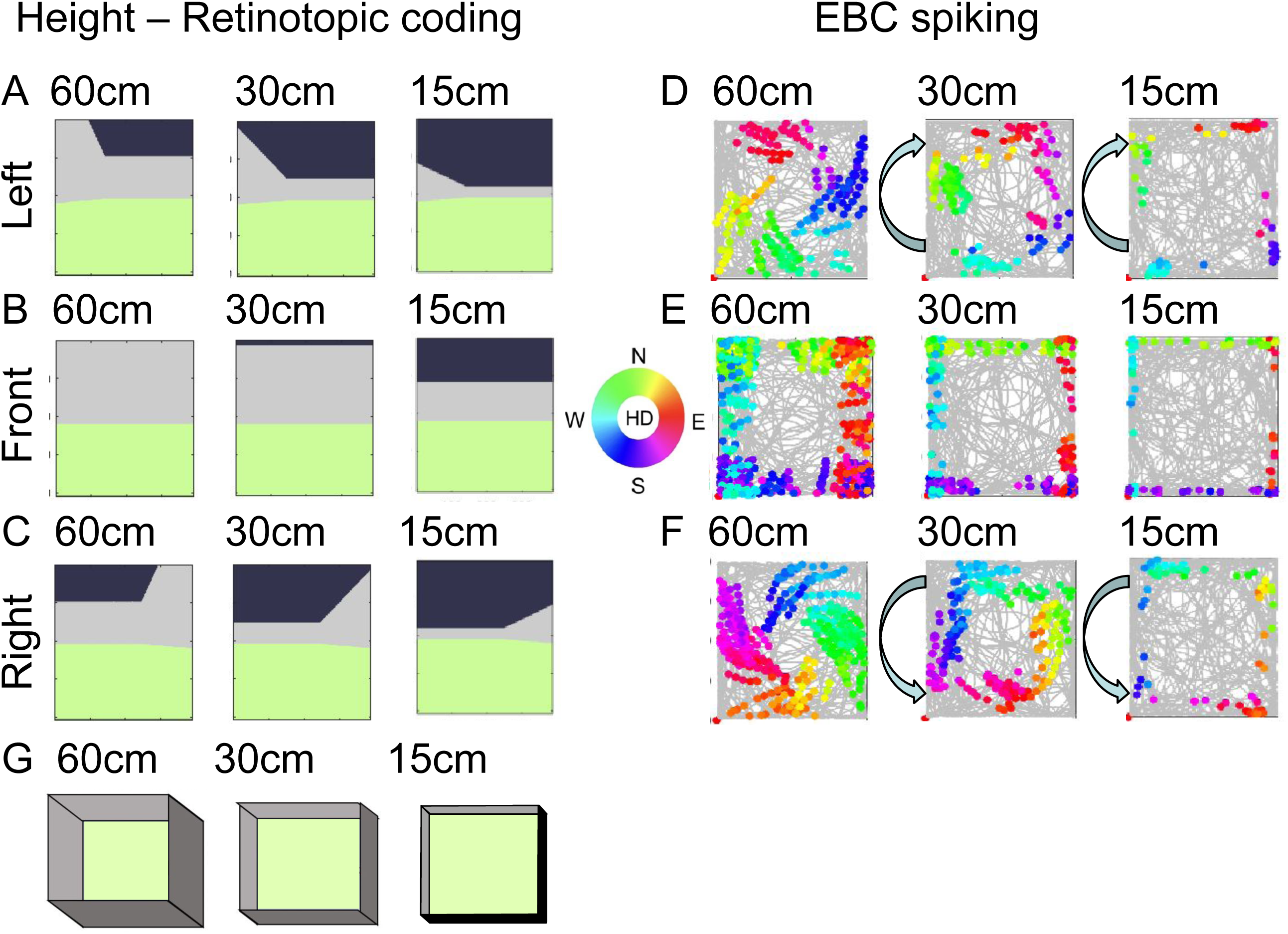
Firing properties of neurons change when boundary walls are higher during sensory input (without changing the memory weights). A. Retinotopic inputs for position at 12cm from left wall at three heights 60cm, 30cm and 15cm. (There is no response at lower wall heights such as 4 cm or 0 cm). B. Retinotopic inputs for 12 cm from wall in front at three heights. C. Retinotopic inputs for 12 cm from right wall at three heights. D. Spiking response for model EBC with left wall memory shows shift in clockwise direction for lower walls (30cm, 15cm). E. Spiking of model EBC with memory for wall in front shows response at greater distance for tall wall (60cm), and closer distance for shorter walls (30cm, 15cm). F. Spiking model EBC with right wall memory showing spiking response that shifts counterclockwise for lower walls (30cm, 15cm). See text for more description of these shifts. G. Schematic showing the three different wall heights tested in these simulations.

Figure 7D shows the spiking response generated by the matching of the 60 cm sensory template for a left side wall with the sensory template for all visual input for 60 cm walls, 30 cm walls, or 15 cm walls. This change in sensory input without a change in memory weights is why the spiking shifts in the different examples in different columns shown in Figure 7D.

This description shows changes in spiking response observed when the height is decreased from Normal to Lower during the sensory input during foraging. For example, in Figure 7D, column 30 cm, the spikes continue to respond to a wall on the left side of the animal (based on the view during memory being of a wall on the left). However, the response shifts to occur at a closer distance from the wall in front of the animal, because the matching of a lower wall with a normal wall memory will occur at a closer distance (a closer low wall looks similar in height to a more distant high wall). Because the spiking depends on both the wall to the side and the wall in front of the animal, the spiking fields show a clockwise rotation, because the matching to the memory occurs closer to the wall in front of the animal when the wall is on the left. For example, the red spikes (for eastward movement) by the north wall shift to the right, and the purple spikes by the east wall shift downward.

The same shift affects spiking in response to a memory template for a wall directly in front of the animal (Figure 7B). In this case, the current sensory input at 12 cm from the wall in front of the animal shows a shorter wall (Figure 7 column 30 cm and 15 cm). In Figure 7E, as the animal forages in the environment, the shorter sensory input (Figure 7E, 30 cm and 15 cm) matches at a closer distance to the wall. This causes spiking to shift to being closer to the wall. For example, the red spikes for moving eastward toward the east wall occur at a shorter distance from the east wall.

Similarly, the spiking plots in Figure 7F for an EBC responding to a wall on the right side also occur at a closer distance from the wall compared to the normal wall sensory input in Figure 7C. However, in contrast to the left side EBC, the spiking fields for the right side EBC show a counterclockwise rotation, because the matching to the memory of a wall in front of the animal occurs at a closer distance when near the right side wall in Figure 7F. The shift is counterclockwise instead of clockwise because the red spikes against the south wall shift to the right in Figure 7F, in contrast to red spikes against the north wall in Figure 7D. The counterclockwise shift also occurs because purple spikes that shift downward do this against the west wall instead of against the east wall. Thus, the retinotopic memory weights predict a rotation in different directions for different walls.

### Head-centered model response to height

Figure 8 shows examples of EBC responses generated with memory features at the center of each plot, based on synaptic weights modified in response to features of a left-side barrier in head-centered coordinates shown in Figure 8A. Note that these memory features in Figure 8A are used for all the simulation outputs in different rows in Figure 8B. Figure 8A also shows the head-centered array from the full sensory input of the surrounding wall of the environment at a specific time step for the 60 cm wall (activating all levels of vertical representation), as well as for 30 cm (surrounding wall appears in half as many levels) and 15 cm (showing only one level of activation of the internal array). Note that the memory template at the center of each array in Figure 8A maintains the same height, coding the wall segment on the left at all vertical heights.

**Figure 8.**
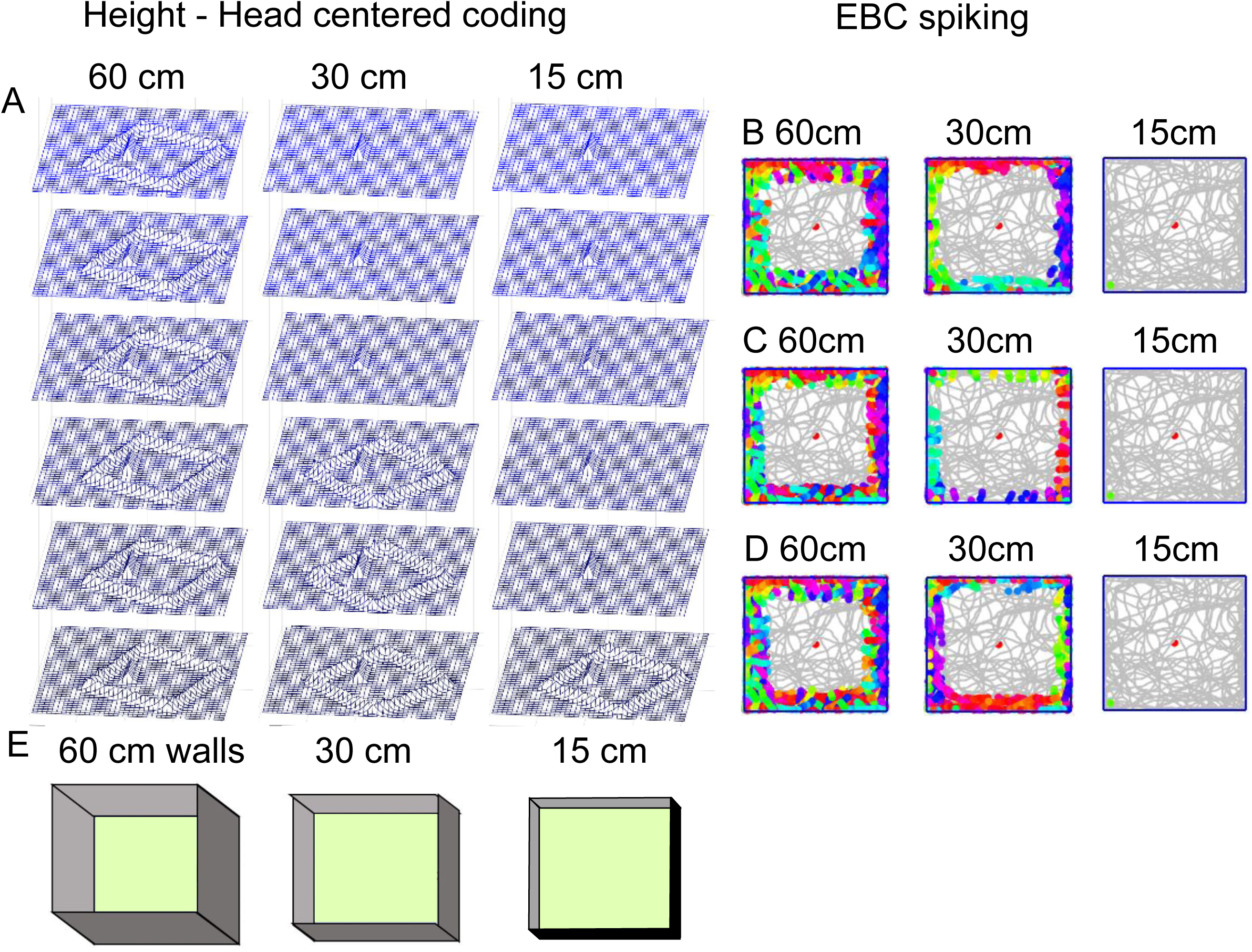
Modeled spiking response during change in boundary wall heights for head-centered neural arrays. A. Head-centered neural arrays representing input during matching for different boundary heights in columns 60cm, 30cm, 15cm. The bump that appears at all levels in center represents synaptic weights encoding memory of a 60 cm wall segment nearby on the left. B. Spiking of model EBC with 60 cm left wall memory for sensory input during matching at 60cm, 30cm and 15cm wall heights. C. Spiking of EBC with front wall memory. D. Spiking of EBC with right wall memory. E. Schematic illustrating the three different wall heights (60 cm, 30cm, 15 cm) simulated during matching in this figure.

The first row of EBC spiking (Figure 8B) demonstrates the spiking response for the left side template shown in Figure 8A. This template generates a strong matching signal for sensory input of walls on the left side of the animal in Figure 8B. The template matching with sensory input generates a weaker response for the smaller vertical extent of sensory input in the 30 cm column of Figure 8B. The template matching with sensory input generates a very weak matching response for the smallest vertical extent of 15 cm in Figure 8B. This falls below threshold and results in no spiking response in Figure 8B, 15 cm.

A similar pattern appears in Figure 8C for head-centered templates coding a wall segment in front of the animal (template not shown), and in Figure 8D for head-centered templates coding a wall segment to the right of the animal (template not shown). In both of these cases, there is a strong match for the sensory input at full height (60 cm), and there is a weaker response when the sensory input contains a shorter wall (30 cm). In both of these cases, the weakest response occurs when the sensory input contains the shortest wall (15 cm), in which case the matching process falls below threshold and there is no spiking response to the walls. Thus, the head-centered simulations predict a change in magnitude of firing rate for decreases in height of the wall, but these simulations do not predict the clockwise or counterclockwise shift of firing because the three dimensional internal representation is not subject to the shift in location of matching that occurs when matching a retinotopic memory template of a 60 cm wall with sensory input of a shorter wall.

### Simulations of EBC firing to changes in barrier tilt

The head-centered and retinotopic models were also tested for their response to different vertical tilts of a barrier that was positioned to have its base stretching across the entire center of the environment in the north-south orientation. In simulations of subsequent foraging periods, this additional barrier was changed to different degrees of vertical tilt so that it blocked movement into one half of the environment as the vertical tilt increased from zero. The simulations started with a 0 degree tilt allowing foraging throughout a 1 meter square environment. The next foraging period was with a 30 degree tilt of the central barrier that blocked movement into one half of the environment, but would have different x, y coordinates for the base, middle and top of the wall. Simulated foraging was also tested with a 60 degree tilt of the central barrier and a 90 degree tilt equivalent to the surround boundaries.

This environmental manipulation was tested in simulations using the head-centered representation in which the neural array would code the difference in x, y position for different vertical coordinates long the barrier (z dimension). For example, for the 30 degree tilt, this would have the neural code of the base with a barrier at the center of the environment, but for higher levels the neural code would represent the barrier at greater distances from the animal along the tilt of the wall. This resulted in a lower match than the vertical wall, and resulted in lower firing rates along the tilted wall for 30 degrees and 60 degrees, as visible in Figure 9Aii and 9Aiii, which have fewer purple spikes along the tilted wall as the animal is running south with the tilted wall on its left. Similarly, in Figures 9Bii and 9Biii, there are also fewer green spikes along the tilted wall as the animal is running north with the tilted wall on its right. The vertical inserted wall in Figure 9Biv resulted in a match of sensory input and head-centered template with firing comparable to the vertical walls in other parts of the environment.

**Figure 9.**
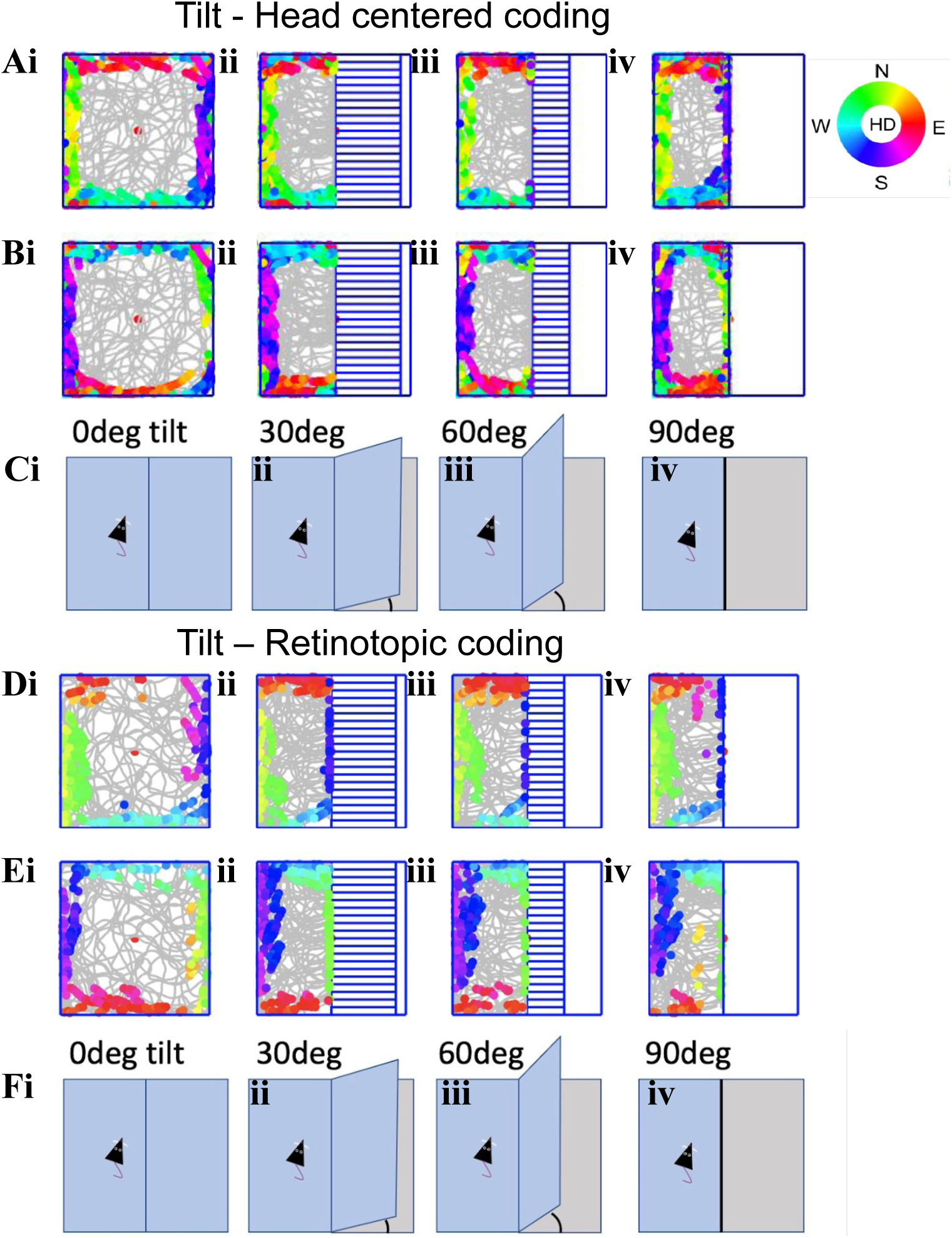
Modeling barrier tilt effects on EBC responses. Ai-Aiv. Head-centered model of spiking responses for left side EBC template in open field (i), with 30 degree tilted barrier (horizontal blue lines), (ii), 60 degree tilted barrier (iii) and 90 degree vertical barrier (iv). Note lack of spiking response to 30 and 60 degree tilted barriers (ii, iii). Bi-Biv. Head-centered model spiking response for right side EBC. Ci-Civ. Schematic showing the different levels of tilt of the central barrier in the environment. Di-Div. Retinotopic model of spiking for left side EBC in open field (i), 30 degree (ii), 60 degree (iii) and 90 degree vertical (iv). Spiking occurs near tilted barrier (blue spikes) and shows extra spiking (violet) for 90 degree barrier (iv). Ei-Eiv. Spiking of retinotopic right side EBC shows simulated spiking near the tilted barriers (green).

The retinotopic simulation of the same paradigm had some differences in response. In this case, the retinotopic view of the inserted wall seems to better match the stored retinotopic image of a 90 degree wall, so there is still some residual firing to the 30 degree and 60 degree tilted wall. This is visible as some residual purple dots in Figures 9Dii and 9Diii, showing spiking along the center tilted wall as the animal runs south with the tilted wall on the left, and in Figures 9Eii and 9Eiii there are some green spikes along the tilted wall as the simulated animal runs north with the tilted wall on its right. In addition, the 90 degree inserted wall shows fewer purple dots (spikes) in Figure 9Div compared to the purple dot spikes for the 90 degree east boundary wall in Figure Di. This is because the retinotopic image has a lower match by the 90 degree inserted wall because the current sensory retinotopic image includes only half as much of the view of the south wall compared to the template learned with a wall on the left. Similarly, Figure 9Eiv shows fewer green spikes to the 90 degree inserted wall compared to the spiking to the east boundary wall in Figure 9Ei. Thus, there are differences in the nature of the spiking activity induced by the matching to memory template when the template is head-centered (Figures 9A and 9B) versus retinotopic (Figures 9D and 9E).

### Simulations of EBC firing to changes in barrier orientation

Testing different models can also benefit from analyzing the response to inserted barriers at different horizontal orientations. The simulations presented in this section show how memory weights for both the head-centered features and retinotopic feature responses can be used to predict EBC responses for different orientations of barrier walls inserted into the environment.

#### Retinotopic simulations of barrier orientation

Figure 10A and 10B shows examples of EBC responses to inserted barrier walls at different horizontal orientations, using memory weights coding visual features in retinotopic coordinates from the starting images shown in Figure 10Ai and 10Bi. The subsequent panels in each figure row show the spiking response based on the dot product of current retinotopic sensory input features with the memory weights of the retinotopic features based on the view in column i. The response in an open field is shown in Figure 10Aii and 10Bii. The response to a barrier inserted with the long axis along the East-west dimension is shown in Figures 10Aiii and 10Biii. The response to a barrier inserted with the long axis in the North-south dimension is shown in Figure 10Aiv and 10Biv. The response to a barrier inserted diagonally into the environment is shown in Figure 10Av and 10Bv, and then another open field example is shown in Figure 10Avi and 10Bvi.

**Figure 10.**
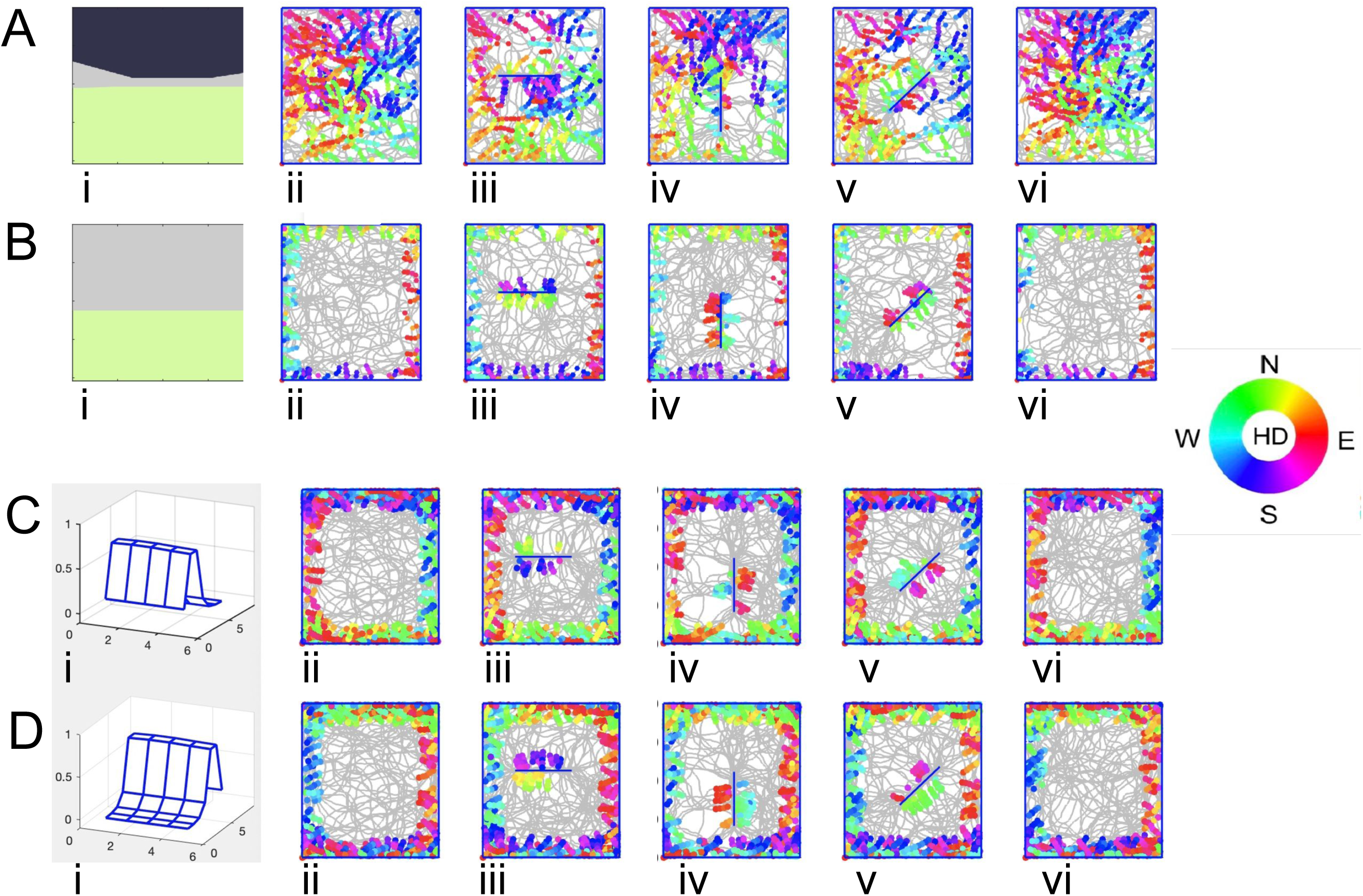
Models of EBC responses for barrier horizontal orientation. Left column shows stored templates. Subsequent columns show gray trajectory show similar EBC spiking response in an open field (column ii) and to 3 inserted barrier orientations (columns iii-v), then another open field (column vi). A. Simulated encoding of a retinotopic memory array for distant barriers when the barrier is behind animal (10Ai). The subsequent matching with retinotopic sensory input generates an EBC spiking response (10Aii-vi) that is not restricted to the boundary and is altered by the inserted barrier. B. Simulated encoding of a retinotopic memory for a wall close in front (10Bi) shows better match of EBC responses (10Bii-vi) for a barrier in front of the animal. C. Simulated encoding of the head-centered memory array for barrier behind animal (10Ci) and five subsequent simulations testing matching of head-centered egocentric input in different sessions (10Cii-vi). This shows more consistent and localized spiking responses to a barrier behind the animal. D. Head-centered memory for barrier in front of animal (10Di) shows clear EBC spiking response to walls in front of animal (10Dii-vi).

Note that the simulation can effectively account for responses to boundaries and barriers in front of the animal as shown in the row in Figure 10B. This works because the retinotopic input primarily contains a segment of the floor and a segment of the wall, and this particular configuration of features (the color elements and the boundary) only match in locations when the animal is directly facing the boundary in the plots. In addition, the view is relatively similar when the animal faces the inserted barrier in Figures 10Biii, 10Biv and 10Bv, and because it codes a view close to the barrier it is not affected by the relationship of the barrier to the surrounding environment (though this barrier/surround relationship will alter firing for examples when the viewpoint is at a greater distance from the wall).

In contrast, the spiking response of an EBC using retinotopic input to generate a response going away from the wall is not very localized to the wall (Figures 10Aii-10Avi). This is partly because the retinotopic view (Figure 10Ai) when going away from the wall is dominated by a very large ground plane (green) and the distant view beyond the walls (black), with only a small set of wall features appearing in gray across the center of the retinotopic view. This results in difficulties in discrimination of dot product values for different parts of the environment. If the dot product threshold is set too high, there will be no response anywhere. If the threshold is set lower, the dot product crosses it in a broad variety of locations as shown for the open field foraging trajectory Figure 10Aii. Note also that the insertion of the barrier in the center of the environment strongly changes the image when looking away from the surrounding walls. This results in a reduction of the spiking activity near the north and south boundary wall for an East-west inserted barrier as shown in Figure 10Aiii, when compared to the more extensive spiking in the open field in Figures 10Aii and 10Avi. This also results in a reduction in spiking activity for the East and West boundary walls when a North-south barrier is inserted (Figure 10Aiv), and reductions of spiking near all walls when a diagonal barrier is inserted (Figure 10Av). Thus, the retinotopic framework predicts that boundary responses in an open field will show changes when a barrier is inserted, and these changes will depend on the relative orientation of a boundary wall relative to the inserted barrier.

### Head-centered simulations of barrier orientation

Figure 10C and Figure 10D show examples of EBC responses generated with memory features based on synaptic weights modified in response to features in head-centered coordinates shown in Figures 10Ci and 9Di. As in Figures 10A and 10B, the subsequent panels in these rows show the spiking response based on the dot product of current head-centered sensory input features with the memory weights of the head-centered features based on the view in the column in Figure 10Ci and 9Di. The response in an open field is shown in Figure 10Cii and 10Dii. The response to different orientations of inserted barrier are shown in the subsequent panels in Figs. 10Ciii-v and 10Diii-v.

In contrast to the retinotopic image, the head-centered representation shows more consistent spiking responses to barriers that are both behind (Figure 10C) and in front (Figure 10D) of the animal. In Figure 10C, the local head-centered array used to train the synaptic memory weights is centered behind the animal. (Note that extent of the array shown in Figure 10C is +/- two units in the front-back dimension and +/- 3 units in the left-right dimension relative to the coordinate origin centered on the animal in egocentric space). When the full head-centered sensory input array interacts with the memory synaptic weights, the neuron receiving this input (computing the dot product) crosses threshold when the boundaries are behind the animal in the open field as shown in Figure 10Cii. This spatially restricted response to a barrier behind the animal is a benefit of a head-centered representation versus a retinotopic representation.

You might ask how does the sensory input represent a barrier behind the animal? This is because the head-centered representation in the model is simulated to undergo updating of the feature array based not only on visual input but on current velocity on each step (including both translation and rotation). Thus, if the animal turns, its internal representation of the sensory input array is updated to include features in head-centered coordinates behind the animal that are no longer visible in the visual field. You might also ask why this isn’t done for the retinotopic representation. This is because the transformation of a retinotopic image cannot be done with a simple linear matrix multiplication which can be done using a linear affine matrix for the head-centered array. The standard way to update a retinotopic image is to transform it back to an egocentric head-centered representation, then perform the affine rotation or translation with a matrix multiplication, and then recompute the projection transform. The bottom line is that head-centered representations allow for more efficient updating of the array.

As seen in the figure, the neuron crosses threshold also when the inserted east-west barrier is directly behind the animal in Figure 10Ciii. Note that the firing continues to be more uniform for the north boundary of the environment in Figure 10Ciii. This appears as blue and violet dots indicating southward head direction just below the North boundary, and to the south of the inserted barrier. This similar firing to boundaries is because the local head-centered array only detects the north boundary behind the animal and is not influenced by the inserted barrier until that barrier is behind the animal (i.e. when the animal is heading south from the barrier (blue dots below inserted barrier). The same thing can be seen for the inserted north-south barrier (e.g. red dots indicating eastward movement away from barrier) and for the inserted diagonal barrier (with violet dots indicating south-eastward movement away from the barrier. Thus, the synaptic weights created by an array of head-centered features is more effective for simulating responses to a boundary or barrier behind the animal. The example in Figure 10D shows similar behavior for an array created based on a local feature array with the boundary wall in front of the animal (Figure 10Di) and generates spiking when the animal is headed toward the boundary or barrier (i.e. blue dots north of the south wall in Figure 10Dii and north of the barrier in Figure 10Diii). Thus, there appear to be advantages of the head-centered representation over the retinotopic representation when simulating responses to barriers inserted into the environment at different horizontal orientations.

### Head-centered responses dependent on current velocity

The neural representation of barriers might be important for computing trajectories that avoid barriers, as was used in previous simulations of the computation of goal-directed trajectories (Erdem & Hasselmo 2012, Erdem & Hasselmo 2014). If barrier codes are being used for obstacle avoidance, then the salience of a barrier might depend upon the current velocity of an animal. For example, the neural response to a barrier in front of an animal might be much more behaviorally important if the animal is moving toward the barrier at 90 cm/sec versus 2 cm/sec. Thus, we were interested in simulating how the egocentric neural response to a barrier might depend upon the current velocity of the animal. (Similarly, a barrier might be more important in front of the animal than behind the animal, though detection of a barrier behind the animal could be important as a potential source of protection from predators).

To perform these simulations of the effect of running speed on egocentric coding of barriers, we used the ray tracing algorithm that we used to create the foraging trajectory of the animals. This forward scanning is inspired by previous models using forward scanning for navigation (Erdem & Hasselmo 2012, Erdem & Hasselmo 2014), and by experimental data indicating that forward scans may be generated by grid cell activity in medial entorhinal cortex (Vollan et al 2024). In initial simulations, this ray tracing algorithm projects a vector forward from the animal and detects whether the animal will run into a barrier within a thresholded distance (Figure 11A2). This thresholded response to the intersection of a ray and a barrier wall already provides a simple way to represent a possible mechanism for the egocentric response to walls. We modified this obstacle avoidance algorithm further by testing different directions of ray tracing forward from the animal (e.g. to model a response to a barrier on the left side of the animal, this would involve ray tracing at a 90 degree angle from the current head direction of the animal). In addition, to test the effect of running speed, we made the length of the projected ray dependent on running speed, and then computed the intersection based on time of intersection (collision) rather than based on the distance of intersection. The effects of this manipulation on simulations of egocentric boundary coding are shown for egocentric coding of wall distance relative to the animal in Figure 11. The figure shows that at low running speeds with short forward trajectory scanning, the network shows a mean egocentric response to barriers close to the animal (Figure 11A1). This corresponds to trajectory plots in which the scans are shorter and the spiking occurs closer to the walls (Figure 11A2). For faster running speeds with longer forward scans, the egocentric response to barriers occurs at greater distances in front of the animal (Figure 11B1 and Figure 11C1). These simulations correspond to trajectory plots with longer scans (blue lines) and with spiking that can occur at greater distances from the walls (Figures 11B2 and 11C2).

**Figure 11.**
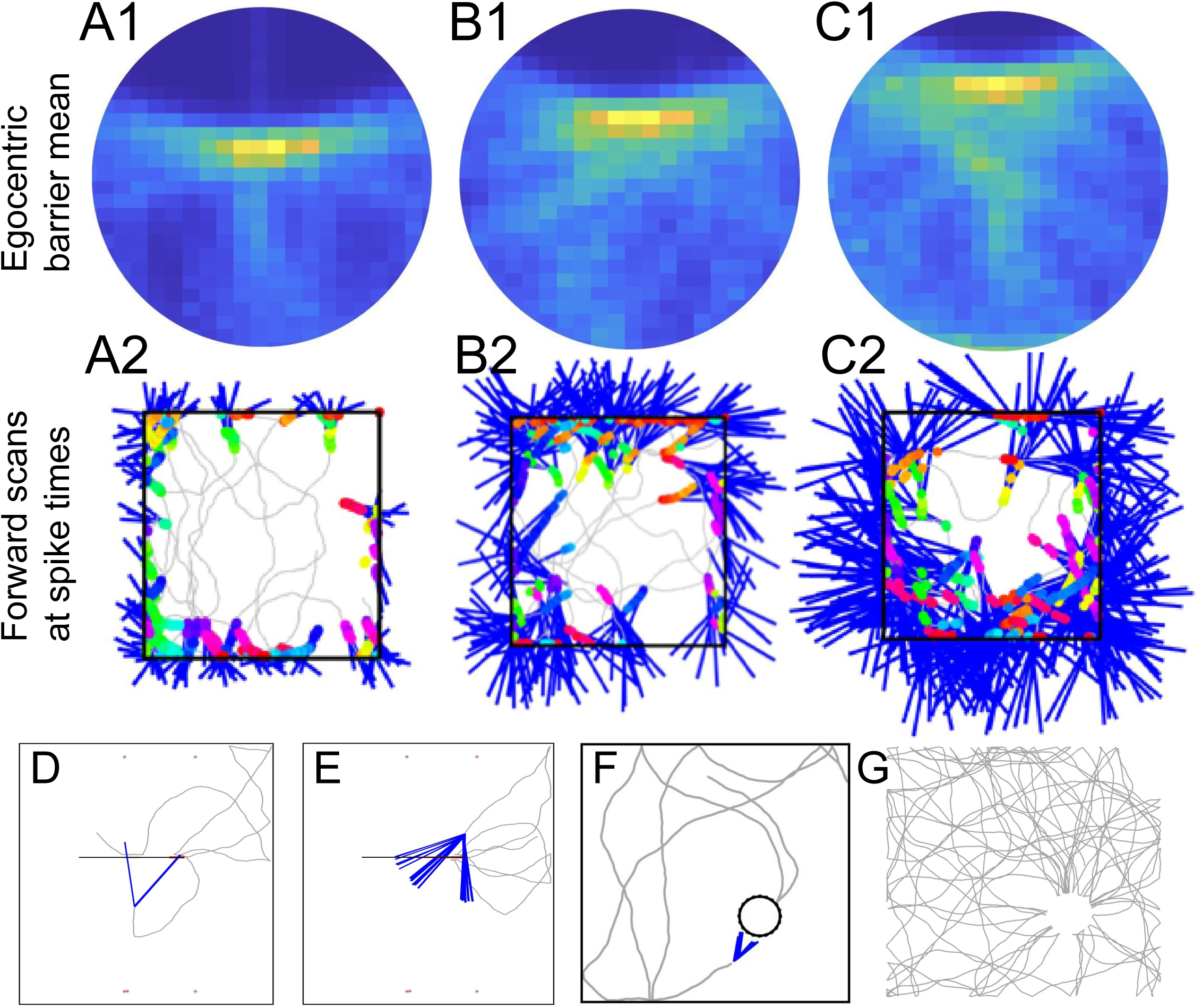
Models showing egocentric coding with forward planning scaled to different running speeds. Egocentric plots of barrier location during simulated spiking are shown in top row, and simulated forward planning scans are shown in bottom row (blue lines are forward scans from spiking locations at colored dots). A1. At lower running speeds, the mean egocentric barrier location for spikes is closer to the center (agent location) because forward planning (A2) predicts collision with head-centered code of barrier at shorter distances. B1, C1. At higher running speeds, the egocentric plot of barrier location during spiking is more distant. This is because the longer forward planning trajectories (B2, C2) with higher running speeds results in firing of EBCs at a greater distance from the barrier. D. Example showing a foraging trajectory (gray) and focusing on only one brief period of forward scanning (blue lines) at 30 degree angles to the left and right of animal to detect the central barrier. E. Another example of detection of the central barrier by forward scanning during a short period of time. F. Forward scanning to detect a circular inserted novel object. G. Trajectory guided by forward scanning that guides attraction to the novel object showing movements converging on object which corresponds to attraction to novel objects in animals well habituated to the environment.

The process of forward scanning is further illustrated in Figure 11D-G. Rather than showing a summary after an extended period of time, Figures 11D and E show just a few forward scans when the animal is near the central barrier in the environment. In these examples, the simulation is generating forward scans at 30 degree angles alternating to the left and right of the animal, consistent with recent data from the Moser lab (Vollan et al 2024). Forward scans can be used to generate the position of egocentric boundary cell spiking as shown in Figure 11A and B, which could allow obstacle avoidance at short distances. These forward scans can also be used by the animal to detect novel barrier features and to be attracted to these novel features, as shown in Figure 11D and E. This attraction to a novel barrier was used in Figure 10 to accurately reflect the fact that animals sample the central barrier more than would be expected by random movement. The attraction to an inserted circular novel object is shown in Figure 11F and G, where the trajectory visits the central object more frequently than expected by chance.

### Egocentric responses in darkness

We also simulated the egocentric response to barriers in darkness. This immediately raises important issues for the response of animals using retinotopic or head-centered coordinates of visual features, as the sensory input would be absent when the animal navigates in complete darkness. In this case, it is necessary to update the head-centered coordinates or ray tracing mechanism based on sensory input about barrier position such as a previous collision with a barrier.

We used a framework in which the animal’s current estimate of its position in the environment would be reset each time it touched a barrier, and then the update would depend upon the path integration of running velocity (speed and direction) as the ongoing integral of running velocity will estimate current position. This allowed simulations of egocentric responses to barriers in darkness based on the path integration of current location, which was also used for updating head-centered representations of the barriers. This was combined with the ray tracing extrapolation of current trajectory to determine whether there was an impending collision with a barrier, and this would drive the spiking of egocentric boundary cells.

As shown in Figure 12, the combination of path integration with ray tracing allowed simulation of egocentric boundary response to barrier position in darkness. As shown in Figure 12Ai, Bi, Ci and Di, the direction of ray tracing could determine the egocentric coding direction. Figure 12 Ai shows how ray tracing at a 90 degree angle to the left of the current head direction could simulate an EBC responding to the left side of the animal (left EBC). Figure 12Bi shows that ray tracing at a 90 degree angle to the right could simulate a right EBC. Figure 12Ci shows that a ray going directly behind the animal (180 degree angle) can simulate an EBC responding to a wall behind the animal. Finally, the standard forward ray tracing used for obstacle avoidance (0 degree angle) can simulate the response to a boundary directly in front of the animal.

**Figure 12.**
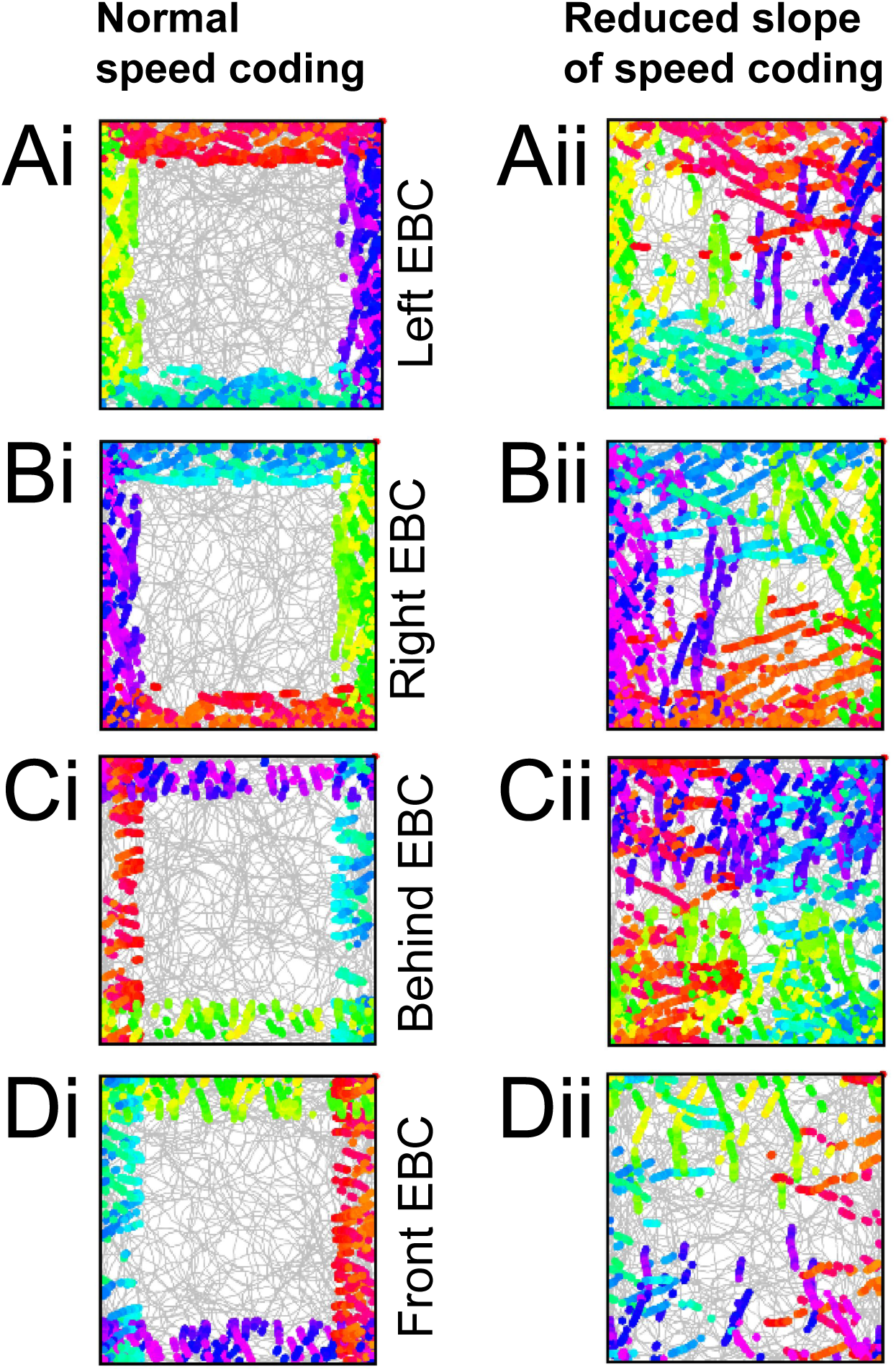
Simulations performed in darkness so that the head-centered array must be updated based on the estimate of current location. The forward planning model uses forward sequences based on current velocity that generate spiking if neural activity interacts with barrier memory within a neural time interval. The left column shows simulations that represent EBC spiking responses with effective path integration based on current location and forward trajectory sampling in different directions: Ai, sampling to the left side (Ai) detects a barrier to the left, Bi, sampling to the right side detects a barrier on the right, Ci, scanning behind the animal, as also shown in some cases by (Vollan et al 2024) detects a wall behind the animal, Di scanning to the front of the animal detects wall in front of animal. The right hand column shows distortions of EBC spiking activity predicted for impaired speed coding (reduction in the slope of firing relative to running speed) due to inactivation of basal forebrain GABAergic input. The effect of reducing the slope of speed coding impairs path integration and results in spreading of spiking responses for all cells including the left side EBC (Aii), right side EBC (Bii), back EBC (Cii), and front EBC (Dii).

This internal representation of barriers would be very dependent on accurate path integration. Anything that seriously distorts the coding of speed should alter the accuracy of egocentric boundary cell firing in darkness. In particular, it has been shown that inactivation of GABAergic neurons appears to decrease the slope of the tuning curve of neuronal firing rate relative to running speed (Robinson et al 2023). This change would be expected to alter the updating of position estimates by path integration, and could result in a distortion of egocentric boundary cell firing due to inaccurate estimates of distance during the period since the previous encounter with a boundary. This would stretch out responses in many cases due to the animal thinking that it is still within the range of a barrier even though it has actually moved out of range of the barrier. Simulation of a 60% decrease in slope of firing rate to running speed causes the distortion of egocentric boundary cell firing shown in Figure 12Aii, Bii, Cii, and Dii. In each case the position estimate becomes less accurate and the firing in response to estimated position of boundaries becomes larger because the simulated circuit has underestimated the amount of movement away from the boundaries.

## DISCUSSION

The models presented here show that egocentric boundary cell responses during open field foraging (Alexander et al 2020a, Alexander et al 2020b, Alexander et al 2023, LaChance & Hasselmo 2024, Malmberg et al 2023, Malmberg et al 2024, van Wijngaarden et al 2020) can be simulated using sensory input arrays in either retinotopic or head-centered. However, the simulations of these different coordinate systems make different predictions about the response of egocentric boundary cells to manipulations of the environment such as changes in the height of environmental boundaries, or changes in the vertical tilt or the horizontal orientation of inserted barriers in the environment.

The framework presented here used a standard approach to neural modeling in which memory weight matrices are formed by Hebbian modification of associations (Kohonen 1984, McNaughton & Morris 1987). The framework presented here formed associations during an encoding process prior to foraging during which Hebbian modification strengthens synapses between presynaptic sensory input neuron arrays in visual cortices activated by visual input and individual postsynaptic retrosplenial neurons activated at an array of locations and head directions. After encoding, during subsequent foraging, the sensory input causes retrieval that spreads across the weight matrix, as simulated by the dot product of the input array with each row of the weight matrix. This determines which postsynaptic retrosplenial neuron has a sufficient match (dot product) to cross threshold and generate a spike at that time point of the trajectory corresponding to a specific location and head direction. These differences in dynamics between encoding and retrieval could be regulated by cholinergic modulation during encoding (Hasselmo 1993, Hasselmo 2006), or by different phases of theta rhythm oscillations (Hasselmo et al 2002, Hasselmo & Stern 2014).

For open field simulations, realistic neuronal responses can be generated for egocentric boundary cells coding walls to the left or right of the animal or in front of the animal. In the open field, this works for sensory input arrays using either retinotopic or head-centered coordinates. Retinotopic coordinates use the horizontal and vertical positions of features projected onto the image plane, whereas head-centered coordinates code the distance and angle of environmental features in 3-dimensional head-centered coordinates (changed to front-back, left-right and up-down coordinates). Further simulations show that manipulations of barrier properties such as height, vertical tilt and horizontal orientation will alter the neural response of the simulated cells in different manners dependent on the use of head-centered or retinotopic coordinates. Previous simulations have shown how sparse encoding and matching of retinotopic images can generate egocentric boundary cells (Lian et al 2023), but have not yet explored the effect of manipulations of barrier parameters.

As shown in Figure 7, for a retinotopic model, the changes in boundary height predict a clockwise or counterclockwise rotation of the neural firing pattern of neurons tuned to left or right side walls, and predict a change in the width of firing fields with height of walls. In contrast, Figure 8 shows that the change in the height for the head-centered model only causes a change in the firing rate of egocentric boundary cells and not location of firing. As shown in Figure 10A and B, for retinotopic models, the response of neurons coding walls behind the animal should change in response to the angle of inserted barriers, with east-west wall boundary wall responses affected by north-south inserted barriers, whereas north-south boundary wall responses would be affected by east-west inserted barriers. In contrast, the head-centered model shown in Figure 10C and D shows more consistent responses to barriers in front and behind the animal despite the insertion of barriers at different orientations.

Ultimately these egocentric representations must influence the formation of allocentric representations, though allocentric coding by grid cells and place cells were not simulated here. Allocentric coding represents the location of an animal relative to objects and barriers in the environment, and should be useful for the generalization of behavior based on memory coding of goal locations relative to other landmarks when the egocentric positions are changing dramatically from moment to moment. Extensive neurophysiological data has demonstrated the coding of location by grid cells in the entorhinal cortex (Hafting et al 2005) and place cells in the hippocampus (O’Keefe & Dostrovsky 1971) which shift their firing with barrier insertion (Muller & Kubie 1987). Allocentric coding is also used by head direction (HD) cells in RSC and POR (Alexander et al 2020a, LaChance et al 2022, LaChance & Hasselmo 2024). Egocentric coding may also contribute to the generation of cells that respond to the allocentric direction and angle of boundaries (Barry et al 2006, Lever et al 2009, Solstad et al 2008) and objects (Deshmukh & Knierim 2011, Deshmukh & Knierim 2013, GoodSmith et al 2022, Høydal et al 2019). Future work could combine egocentric coding of barriers and running speed to generate allocentric representations of the environment important for goal-directed behavior.

Extensive previous modeling work has focused on the formation of allocentric spatial representations of allocentric boundary cells (Bicanski & Burgess 2018, Byrne et al 2007, Raudies & Hasselmo 2012, Raudies & Hasselmo 2015) and place cells from egocentric boundary cells (Bicanski & Burgess 2018, Byrne et al 2007) usually focusing on two dimensional coding in the azimuth plane. Note that our model focuses on the formation of egocentric boundary cell responses from sensory input during behavior. The simulations presented here do not focus on the later stages of the process modeled in those studies such as the combination of egocentric boundary cells with head direction input to generate allocentric boundary cells (Bicanski & Burgess 2018, Byrne et al 2007). There have also been many previous simulations of grid cell responses based on path integration of movement velocity (Burak & Fiete 2009, Burgess 2008, Bush & Burgess 2014, McNaughton et al 2006, Zilli & Hasselmo 2010). Some models of grid cells address the influences of boundary features on path integration to enhance the accuracy of grid cell firing by having sensory features reset grid cell phase (Alexander et al 2023, Bush & Burgess 2014). However, most previous models of grid cells do not simulate explicit visual sensory input in retinotopic coordinates, though head-centered coordinates have been used to model grid cells as affine transformation matrices (Alexander et al 2023).

Our simulations presented here do not focus on the potential influence of egocentric boundary cells on the allocentric coding by grid cells, but other work in our lab has focused on how models of grid cells could be extended to represent the location of animals, barriers, and objects in the environment (Alexander et al 2023, Hasselmo 2009, Hasselmo et al 2010). Scanning of forward trajectories to detect boundary collisions as simulated in Figure 11 could involve grid cell population codes of location. Forward scanning by grid cells was simulated in early models that used scanning for both obstacle avoidance and goal directed navigation (Erdem & Hasselmo 2012, Erdem & Hasselmo 2014). This model of forward scanning is supported by recent data on decoding of forward scans from grid cell populations (Vollan et al 2024). In related work, the spatiotemporal trajectories coded by grid cells and time cells have been encoded as episodic memories (Hasselmo 2009, Hasselmo 2012) that includes views of barriers and objects at specific locations. This allows forward retrieval of an extended complex spatiotemporal trajectory as an episodic memory (Hasselmo 2009, Hasselmo 2012). Recently, similar model for trajectory retrieval was shown to have a high memory capacity (Chandra et al 2025).

The retinotopic coding used here is very similar to the retinotopic coding used for pixels in the first layer of a convolutional deep neural network (CNN). Rather than training a CNN or other deep neural network, here we maintained neural interpretability by using a single layer for coding the coordinate system transformation into the activity patterns of retrosplenial neurons. Thus, the internal representations used in the simulations presented here are not generated by gradient descent. Many decades of neural network models trained by gradient descent have addressed pattern recognition. For example, deep convolutional neural networks (CNNs) trained by gradient descent have been used extensively for pattern matching in the form of object identification. The first layers of these CNNs match well to simple cells of the primary visual cortex discovered by Hubel and Wiesel. However, somewhat surprisingly, there has not been a clear description of regularities of higher order internal representations that these models generate, partly due to the code being so broadly distributed across multiple large layers, and partly due to the fact that mapping to neural data have usually used transformation matrices that obscure the nature of the representation in the model. We know the nature of neural data in visual cortices that code some object features and individual objects, and we know that the internal representation of the CNNs can be transformed into neural data, but there is no constraint or description of the magnitude or range of CNN representations, and the focus on mean firing rates has ignored the possible role of spike timing or phase coding.

Previous studies have used deep reinforcement learning networks to simulate spatial behavior based on simulated visual inputs and have been shown to generate some neural responses similar to grid cells and place cells (Banino et al 2018, Schaeffer et al 2023, Sorscher et al 2023). A recent model using a more biologically motivated gradient descent model generated spatial representations including head direction cells and egocentric boundary cells (Chapman et al 2024). Further analysis of these models might give insight into the nature of how these spatial neural responses could be generated, but they do not provide an immediate comparison of retinotopic versus head-centered internal representations for the formation of egocentric boundary cell responses.

As described in the Methods section, the sensory input is simulated using standard techniques of computer animation or game engines, in which features are first defined in 3D allocentric coordinates, and the agent movement is simulated. Then, features at each point along the movement trajectory are transformed into egocentric coordinates, and then the projection transform generates retinotopic coordinates. It would be interesting if neurons could work with the definition of features by their coordinates. However, it is not clear how neurons could perform a matching function with only the standard coordinate points of computer vision, as this would require at least three things: 1.) a neural representation of feature coordinates (i.e. neurons that flexibly code a variety of locations), as opposed to the standard neural network approach of using neurons that code retinotopic location and are activated only if the feature corresponds to that location (as in typical simple cell Gabor filtering); 2.) some measure of the relevant density of points, as the coordinates of wall corners are quite sparse; and 3.) solving the problem of knowing which input feature matches with which stored feature. Perhaps neurons could code these coordinates by the phase of spiking relative to network oscillations, or by the intrinsic phase of membrane potential. However, this has not yet been simulated, and therefore the framework presented here utilizes the more standard approach in which individual input feature neurons code the presence of a specific feature at a specific location in the array (the horizontal or vertical position in retinotopic coordinates, or the front-back, left-right, or up-down coordinates for feature position in head-centered coordinates).

In summary, the simulations presented here demonstrate how egocentric boundary cells can be generated according to different hypotheses concerning the role of sensory visual input in either retinotopic or head-centered coordinates, or via forward scanning to detect collisions with barriers. These different hypotheses make different predictions for future experimental tests of the effect of manipulations of environmental features.

## Acknowledgements

This work supported by the National Institute of Mental Health, grant number R01 MH120073. The authors have no conflict of interest.

## Notes

### Competing Interest Statement

The authors have declared no competing interest.

### Summary of Updates

Update to manuscript and figure 1

